# Insights Into Ribosomal DNA (rDNA) Dominance and Magnification Through Characterization of Isogenic Deletion Alleles

**DOI:** 10.1101/2024.03.13.584845

**Authors:** Selina M. Kindelay, Keith A. Maggert

## Abstract

The major loci for the large primary ribosomal RNA genes (35S rRNAs) exist as hundreds to thousands of tandem repeats in all organisms, and dozens to hundreds in Drosophila. The highly repetitive nature of the rDNA makes it intrinsically unstable, and many conditions arise from the reduction or magnification of copy number, but the conditions under which it does so remains unknown. By targeted DNA damage to the rDNA of the Y chromosome, we created and investigated a series of rDNA alleles. We found that complete loss of rDNA leads to lethality after the completion of embryogenesis, blocking larval molting and metamorphosis. We find that the resident retrotransposons – R1 and R2 – are regulated by active rDNA such that reduction in copy number derepresses these elements. Their expression is highest during the early 1^st^ instar, when loss of rDNA is lethal. Regulation of R1 and R2 may be related to their structural arrangement within the rDNA, as we find they are clustered in the flanks of the Nucleolus Organizing Region (NOR; the cytological appearance of the rDNA). We assessed the complex nucleolar dominance relationship between X- and Y-linked rDNA using a Histone H3.3-GFP reporter construct and incorporation at the NOR, and found that dominance is controlled by rDNA copy number as at high multiplicity the Y-linked array is dominant, but at low multiplicity the X-linked array becomes derepressed. Finally, we found that multiple conditions that disrupt nucleolar dominance lead to increased rDNA magnification, suggesting that the phenomena of dominance and magnification are related, and a single mechanism may underly and unify these two longstanding observations in Drosophila.

## Introduction

The ribosomal RNA genes of Drosophila melanogaster are multi-copy, existing as tandem repeats of transcription units for the homozygous 5S rRNA (the 5S array) and the co-transcribed 35S primary transcript comprising the 18S, 5.8S, and 28S rRNAs (the 35S array). The repetition of the rRNA genes is a common feature in eukaryotes with only minor variations (e.g., the 5S and 35S genes are interspersed in a single array in the yeast Saccharomyces cerevisiae) (Bell et al., 1977; Engel and Kobayashi, 2023) Much work on the 35S array – called simply the “ribosomal DNA” (“rDNA”) – has been done in Drosophila, maize, Arabidopsis, yeast, mouse, and human cells (van Sluis et al., 2019; Grummt and Langst, 2013). Reduced rDNA copy number limits translation, leading to a bobbed phenotype in Drosophila (Bridges and Brehme, 1944; Ritossa et al., 1966). bobbed flies express etched cuticles (especially of the dorsal abdomen), reduced length and girth of adult sensory bristles, and developmental delay (Ritossa et al., 1966; Ritossa, 1968). The expressivity of the bobbed phenotype linearly relates to the total number of active rDNA cistrons to a minimum number (Hawley and Tartof, 1985; Terracol, 1986; Paredes and Maggert, 2009a; Tartof, 1973) below which further deletions lead to organismal or cell death. One additional finding of these studies has been that rDNA arrays may contain many more individual 35S cistrons than is strictly necessary for translational capacity, allowing rDNA copy number to vary widely in populations (Lyckegaard and Clarke, 1989; Long and Dawid, 1980).

The repeat structure of the rDNA is intrinsically unstable, contributing to this variation. In yeast, rDNA copies are lost or sequestered in mother cells owing to closed mitoses and, to compensate, the rDNA copy number is regulated by a special process evolved to expand low-copy rDNA arrays back to sufficiency (Koboyashi et al., 1998). The loss and gain of rDNA contribute to the variation observed in populations of individuals. In Drosophila also, copies of the rDNA are lost through development, although the mechanism for excision from the chromosomal array is not well understood (Ritossa, 1968). In Drosophila, the competing process of rDNA magnification can restore bobbed flies to wild-type (Hawley and Tartof, 1985; Ritossa et al., 1966), though it is yet unclear if magnification is an orderly and regulated event, as it is in yeast, or instead is a byproduct of sporadic DNA damage and imprecise repair. Nonetheless, the connections between rDNA copy number changes in yeast as it ages, the potential for a general phenomenon of rDNA loss in aging eukaryotes, and the rapid loss of rDNA in cancer (Xu et al., 2017; Valori et al., 2020; Wang and Lemos, 2017), have led to interest in rDNA copy number dynamics. Drosophila represents an excellent system in which to study loss and magnification of rDNA in part because of the ease in creating otherwise-isogenic rDNA copy number variants (Paredes and Maggert, 2009a). Further, the unusual structure of the rDNA of Drosophila provides compelling insights into the regulation of rDNA transcription itself.

In Drosophila, the non-Long Terminal Repeat retrotransposons, R1 and R2, insert themselves at specific sites within the 28S rRNA genes of some 35S rDNA cistrons (Figure 1A) (Jakubczak et al., 1990). A significant proportion of rDNA cistrons may be so-interrupted by R1, R2, or by both types of insertions in melanogaster, in other Drosophilidae (Malik and Eickbush, 1999) and more generally in the Arthropoda (Jakubczak et al., 1991). The frequencies and distributions of these insertions vary widely between different wild- or laboratory-isolates of Drosophila melanogaster, ranging between 17 and 67% of the X-linked rDNA interrupted by R1s, and 2 to 28% of both the X- and Y-linked rDNA interrupted by R2s (Jakubczak et al., 1992; Wellauer et al., 1978; Long and Dawid, 1979). Thus, only a fraction, as low as 10%, of rDNA may be uninserted in common laboratory strains (Ye and Eickbush, 2006). Nuclear run-on experiments have shown that R1 and R2 are co-transcriptionally expressed with the 28S rRNA as part of the 35S primary rRNA transcript, but are identified and terminated to generate truncated transcripts which are rapidly degraded (Ye and Eickbush, 2006). An additional means of preventing a disruption of 18S–5.8S–28S stoichiometry is the direct transcriptional silencing of R1- and R2-inserted 35S cistrons through sequence-specific or regional regulatory processes (Guerrero and Maggert, 2011; Eickbush et al., 2008; Kindelay and Maggert, 2023) Electron micrographs confirmed that long rDNA primary transcripts – those corresponding to retrotransposon-inserted 35S cistrons – are both under-transcribed and transcribed to varied lengths, indicating absent or incomplete processing compared to the more uniformly-processed uninserted rDNA (Jamrich and Miller, 1984; Chooi, 1979). Correspondingly, expression of R1- and R2-inserted copies has been detected only at very low levels compared to uninserted copies in ovaries, embryos, larvae, pupae, and adult flies of different strains (Long and Dawid, 1979; Kidd and Glover 1981). For these reasons, R1- and R2-inserted rDNA copies are thought to be completely non-functional in terms of ribosome biosynthesis (Terracol and Prud’homme, 1986).

**Figure 1.**
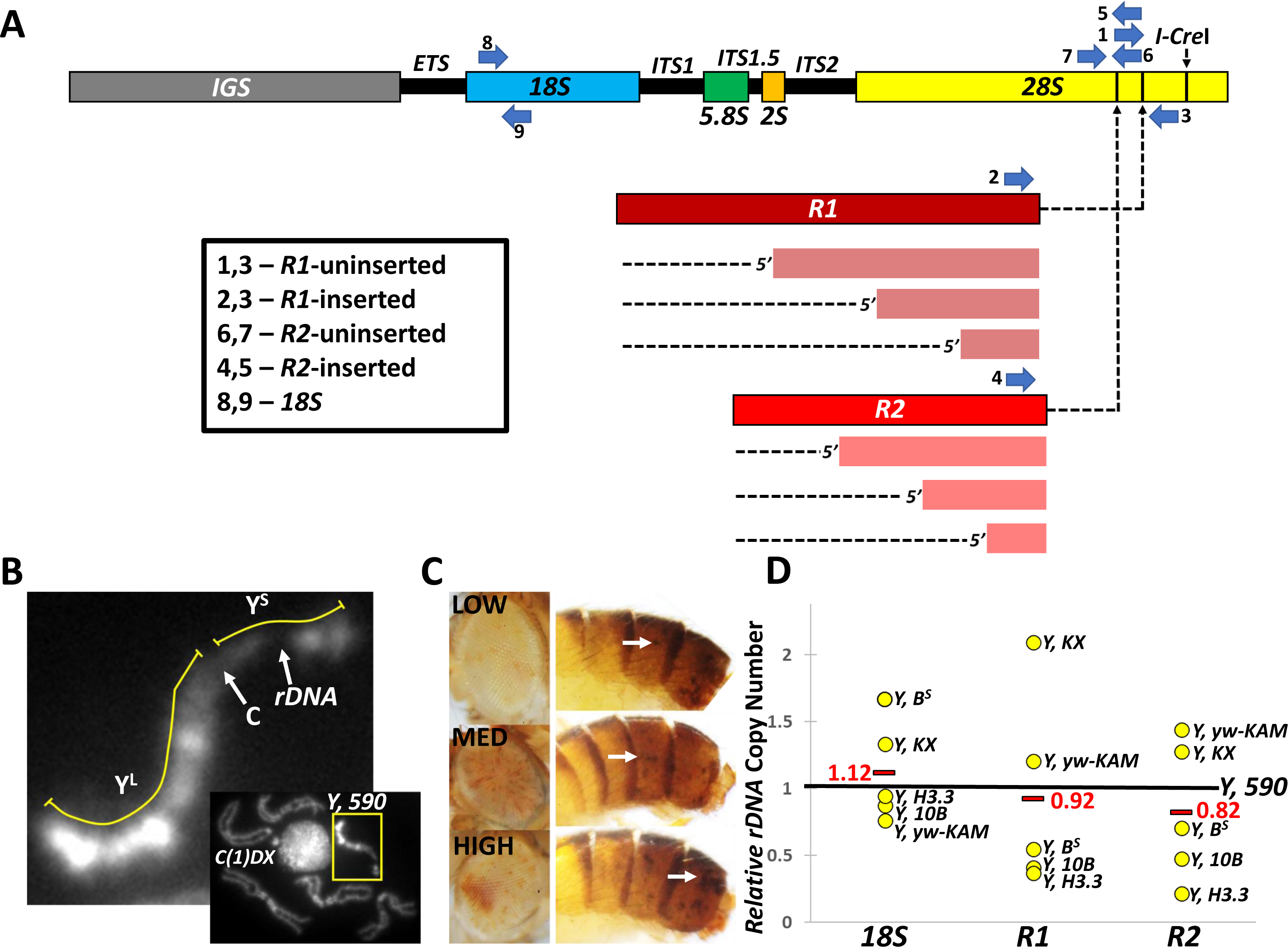
Analysis of *Y*, *590*, *Y* chromosome progenitor for *rDNA* deletion alleles. (A) Schematic of a single ribosomal DNA (rDNA) repeating unit indicating the coding (18S, 5.8S, 2S, 28S) and noncoding regions (IGS is the Inter<u>g</u>enic <u>S</u>pacer, ITS1, ITS1.5, and ITS2 are the Internal <u>T</u>ranscribed <u>S</u>pacers). Red rectangles represent the R1 and R2 retrotransposable elements and dashed arrows indicate their stereotyped insertion sites in the 28S. Orange rectangles and dashed lines represent that not all R1 and R2 elements are full-length; many are 5’ truncations. Numbered blue arrows indicate position of primers used for quantitative PCR (qPCR) or reverse-transcriptase PCR (RT-qPCR) analyses of 18S, 28S, R1, and R2 copy numbers in this study; boxed text indicates which primer sets were used for which class of rDNA. “I-CreI” indicates location of I-CreI meganuclease consensus sequence in the rDNA. (B) Cytology of Y, 590-derived chromosome, including labels of the long arm (Y^L^), short arm (Y^S^), centromeric “primary” constriction (“C”) and rDNA “secondary” constriction (“rDNA”). Genotype of the cell is C(1)DX, y^1^ f^1^ bb^0^/Y, 590^SK11-bb-L^, as apparent from full mitotic chromosome spread (inset); C(1)DX compound-X chromosome is labeled. (C) Examples of Low, Medium and High expression of yellow^+^ and white^+^ due to variegation in the eyes and abdomens of y^1^ w^67c23^/Y, 590 males. See also Supplementary Figure 1. (D) qPCR copy number of 18S, R1, and R2 – relative to Y, 590 – of various laboratory strains. Black bars are the Y, 590 data (set to 1.0), yellow dots are individual data points for indicated chromosomes (see Supplemental Figure 11), and red bars are average of all data (excluding Y, 590).

The bobbed genetics of the rDNA loci is complicated by the internal redundancy of multiple 35S cistrons (with their different retrotranspositional insertion statuses), the redundancy between the two X-linked arrays in females and the X- and Y-linked arrays in males. It has been established that in Drosophila, a dominance relationship exists in that males express only their Y-linked arrays (Greil and Ahmad, 2012). Much remains to be learned about this phenomenon, for example the tissues and cells, and periods of development, in which dominance is observed (Warsinger-Pepe et al., 2020). Additionally, some chromosome combinations escape dominance, and the X- and Y-linked arrays are co-dominant in males (Greil and Ahmad, 2012). The features of the arrays, whether dominance maps to the X or the Y, and how chromosomes may be altered to become dominant are all outstanding questions.

Genetic evidence exists for inter-array compensation as Y-linked arrays with few functional cistrons may be dominant in terms of expression, but recessive in terms of phenotype, depending on the status of the X-linked rDNA array in the genotype. Further evidence for compensation between X- and Y- linked arrays comes from R1 and R2 expression, which have been shown to increase in bobbed mutants as a consequence of increased overall transcription of the rDNA (Terracol, 1986; Long et al., 1981). Despite these behaviors, direct cytological evidence of compensation is lacking.

Knowing how many copies of uninserted rDNA exist in any one individual, how the copy numbers vary in populations, and how expression corresponds to both homologous rDNA copy number and conditions that elicit magnification are critical questions to understand the limits and dynamics of rDNA loss and gain. It is with this in mind that we created rDNA alleles from a natural isogenic Y chromosome that by itself was capable of sustaining the translational demands of the organism. In our studies, we found that the lethal phase of even complete loss of rDNA is late in development, as organisms with no active rDNA complete embryogenesis and hatching, but fail to undergo their first larval molt and die as unnaturally-old 1^st^ instar larvae. The state of rDNA expression corresponds to Histone H3.3 incorporation and the formation of a visible “secondary” constriction in prophase/prometaphase chromosomes, and analysis of these data showed clear evidence for compensatory activation of the X-linked rDNA in what would have otherwise been bobbed flies. Finally, we found that rDNA“ magnifying” conditions corresponded to conditions under which this compensatory mechanism is activated, either by direct manipulation of bobbed alleles or by mutations in genes whose products are necessary for rDNA silencing or chromosome pairing. We propose that dominance inhibits magnification, in which conditions necessary and sufficient for rDNA magnification are those that also elicit X chromosome derepression, unifying rDNA compensation, dominance, silencing, and expression-dependent damage to create the conditions required for unequal sister chromatid exchange-mediated gene magnification.

## Materials and Methods

### Fly strains and genetics

All flies were maintained at 18°C and 70% relative humidity, and crossed or raised at room temperature or 25°C and 80% relative humidity. Strains were maintained and crossed in glass vials on cornmeal molasses agar. Supplemental Figure 11 provides a complete list of genotypes and corresponding stock center numbers or source citations.

### Genetic screen for ribosomal DNA deletions

Heat shock induction of I-CreI meganuclease (Thompson et al., 1992) was performed essentially as previously described (Maggert and Golic, 2005; Paredes and Maggert, 2009a) For each cross, males and females were allowed to freely mate and lay eggs for 2-3 days, then parents were removed. Embryos were allowed to develop to feeding- and wandering-stage 3^rd^ instar larvae. Individual vials were heat- shocked in a circulating water bath at 37°C with fly vial plugs pushed down to prevent larvae from crawling up the vial walls to escape the heat. Time of heat shock and repetition/recovery were varied as outlined in (Supplementary Figure 2).

### 18S, R1, R2 inserted and uninserted copy number determination

Real-Time PCR was performed using the StepOnePlus Real-Time PCR System (Applied Biosystems) with the same cycling parameters as described previously (Paredes and Maggert 2009a, Aldrich and Maggert 2014) To estimate copy numbers of rDNA in bobbed-lethal derivative strains and wild-type laboratory strains, genomic DNA was isolated from 1^st^ instar larvae, 3^rd^ instar larvae, or adults in pools of 10-20 individuals per biological replicate, according to Dobie and colleagues (Dobie et al., 2001) DNA was quantified on a Qubit fluorometer (Thermo Fisher). Each PCR reaction contained Sybr Green 2X Master Mix (Genesee Scientific), appropriate primer pairs (Figure 1A) and gnomic DNA diluted to 2-10 ng/µL per reaction. A master tube of each reaction was split into 4 technical replicates in 96 well plates and run using the comparative “C_T_” method. Primers were: for 18S copy number, d18s.1 5’-AGCCTGAGAAACGGCTACCA-3 ‘and d18s.2 5’-AGCTGGGAGTGGGTAATTTACG-3’; for R2 copy number, YesR2F 5’-ACACACAGTGTTGGCAGACC-3 ‘and YesR2R 5’-TAGATGACGAGGCATTTGGC-3’; for R1 copy number, YesR1F 5’-ATAGGGTGCCGTGGTTGTAA-3 ‘and YesNoR1R 5’-AATTATTCCAAGCCCGTTCC-3’; for uninserted (i.e., neither R1 nor R2) copy number, KR1R2.F - 5’-TTCAAGTAAGCGCGGGTCAAC-3 ‘ and KR1R2.R - 5–’ATTCCAAGCCCGTTCCC-TTG-3’. All data analysis with PCR calculations was performed in Microsoft Excel or Apple Numbers using raw C_T_ values as we have done before (Paredes and Maggert 2009a, Aldrich and Maggert 2014).

### Ribosomal RNA (and *R1* and *R2*) expression

Ten to twenty 1^st^ instar larvae were collected in 1.5 mL microcentrifuge tubes and frozen at -20°C for 15 minutes, then homogenized with plastic pestles in 1 mL Trizol reagent (Ambion). RNA was isolated following the procedure as described (Green and Sambrook 2020). First strand cDNA synthesis was performed with Maxima H Minus Reverse Transcriptase (Thermo Scientific) following the protocol of the kit and using gene specific reverse-complement primers. To detect R2, R2.RT.PG 5’-GCCAACACTGTGTGTGGTCA-3’, to detect uninserted copies, R1R2.RT.PG 5’-CCGAGGTGTAATATCTCCCAC-3’, to detect total copy numbers, 35S.RT.PG -5’-CCGAGGTCTAATATCTCCCAC-3’, to detect R1, R1.R.JA 5’-CCAGCAATCGTATGCTCGCTG-3’; rho1 expression was used as a control, rho1.RT.PG 5’-CTTAGCCGAACACTCCAAATAGG-3’. The cDNA was diluted 1:5 and used directly in Real-Time PCR with the following primers for second strand cDNA synthesis and amplification: for R1, R1.F.PG 5’-GCCTCGTCATCTAATTAGTGACGCGC-3’ and R1.R.PG 5’-CCACGAGCGCAACG-AAAACACG-3’; for R2, R2.F.PG 5’-GGATGTGATGCTCCCGAAAC-3’ and R2.R.PG 5’-CAAGTCCCCGCTTGATT-CGA-3’; for uninserted copies, R1R2.F.PG 5’-GCCTCGTCATCTAATTAGTGACGCGC-3’ and R1R2.R.PG 5’-CCCTTGGCTGTGGTTTCGCTAG-3’; for rho1, rho1.F.PG 5’-GTGGAGCTGGCCTTGTGGG-3’ and rho1.R.PG 5’-CTAGCGAATCGGGTGAATCCACTG-3’, and total rRNA expression, 35S.F.PG - 5’-AGCCTGAGAAACGGCTAC-CA-3’ and 35S.R.PG - 5’-AGCTGGGAGTGGGTAATTTACG-3’.

### DNA Fluorescence *in situ* Hybridization (FISH)

DNA FISH was performed as described previously (Larracuente and Ferree, 2015), 3^rd^ instar crawling larvae were selected and their brains dissected in 1X PBS (Phosphate Buffered Saline, 137 mM Sodium Chloride, 2.7 mM Potassium Chloride, 10 mM Sodium Phosphate dibasic, and 1.8 mM Potassium Phosphate monobasic) in 35 mm x 10 mm cell culture dishes under a SMZ 1500 stereomicroscope (Nikon). The brains were briefly submerged in 50 µL 1X PBT (1X PBS supplemented with 0.1% Tween-20) before incubating in 50 µL 0.5% Sodium Citrate for 10 minutes. Three or four brains were transferred to 20 µL Fix solution (2.5% Paraformaldehyde (pH7.5) in 45% Acetic Acid) on a salinized coverslip (glass treated with Dichlorodimethylsilane and washed with Ethanol, 18 mm or 24 mm) and incubated for 5 minutes at room temperature. Afterward, a Superfrost Plus microscope slide (Fisher) was carefully placed over the coverslip, inverted, and squashed with the firm and experienced thumb of S. Kindelay, then placed on a block of dry ice for 5 minutes or in a -80°C freezer for 15 minutes until frozen. The coverslips were popped off with a razor blade, the slides were incubated in a Coplin jar filled with 100% ethanol at -20°C for 5 minutes, then left to dry at room temperature for an hour.

Once dried, 20 µL of Hybridization buffer (Larracuente and Ferree, 2015) plus 1 µL of probe was pipetted onto the tissue and topped with a coverslip. The slide was incubated in the dark on a heat block at 95°C for 5 minutes. The slide was sealed with stretched Parafilm (Bemis) and incubated at 37°C in a hybridization oven in a humidifying chamber overnight. The next day, the coverslip was gently removed, and the slide washed thrice in 0.1 X SSC (150 mM Sodium Chloride, 15 mM Sodium Citrate) and allowed to dry in the dark at room temperature. A drop of Vectashield (Vector Labs) mounting solution supplemented with 4′,6-diamidino-2-phenylindole (DAPI, 1 ng/µL) was placed over the tissue, secured with a cover slip, sealed with pretty fingernail polish and imaged on a Zeiss Axioskop II mot-plus microscope using a 100X oil objective.

To generate probes, genomic DNA was isolated from a mix of y^1^ w^67c23^ males and females. The genes for the probes were amplified using LongAmp Taq 2X Master Mix (New England Biolabs) with optimized PCR cycling parameters. Primers were: for R1, R1.F 5’-CGGACGTGTTTTCGTTGCGCT-CGT -3 ‘and R1.R 5’-ATGTATGCTTTTCGG-ATCCCTCCGA-3’; for R2, R2.F 5’-TTGGGGATCATGG-GGTATTTGAGA-3 ‘and R2.R 5’-TGATCGC-GGAGGTATGGAAATCT-3’; for 18S, 18s.F 5’-ATT-CTGGTTGATCCTGCCAGTAG-3 ‘and 18s.R 5’-TAATGATCCTTCCGCAGGTTCAC-3’. The PCR products were separated by gel electrophoresis, extracted and purified (Geneclean) and cloned into the pCR4-TOPO-TA vector (Invitrogen). The FISH Tag DNA Multicolor Kit (Invitrogen) was used for nick translation and chemical labelling with Alexafluor (AF) dyes according to the kit instructions, using 1:500 DNAseI and incubating overnight at 15°C. The probes were labelled as follows: 18S probes with AF-488 and R1/R2 probes with either AF-555 or AF-594.

### *Histone H3.3-GFP* Immunofluorescence

Visualization of Histone H3.3 on mitotic chromosomes was done using a protocol adapted from (Larracuente and Ferree, 2015; Greil and Ahmad, 2012). Wandering 3^rd^ instar larvae, either heterozygous or homozygous for the H3.3-GFP transgene, were incubated in a 37°C circulating water bath for 1 hr to induce transient expression of H3.3-GFP, followed by recovery at 25°C for 2 hours. Male larvae were selected and their brains dissected in 1X PBS. The brains were transferred to Sodium Citrate for 5 minutes, then moved to 20 µL of fix solution for 5 minutes on a salinized coverslip. The solution was then carefully pipetted out while avoiding the tissue, and 20 µL of 1X PBT was added and left to incubate for 5 minutes. The PBT was then carefully pipetted out and 20 µL of Fix solution was added. A slide was placed over the coverslip, inverted, and squashed with assured and assertive pressure and placed on a block of dry ice for 5 minutes or placed in a -80°C freezer for 15 minutes until frozen. The cover slips were popped off with a razor blade and washed in a Coplin jar with 1X PBS for 10 minutes and then 1X PBT for 10 minutes. The tissue was covered in 75 µL of Blocking solution (1% Bovine Serum Albumin in 1X PBS) and secured with a square piece of unstretched Parafilm for 1 hour at room temperature in a humid chamber. Excess blocking solution was removed and the primary antibody was added (mouse monoclonal GFP antibody GFP (B-2), sc9996, Santa Cruz) at 1:40 in Blocking solution and incubated in a humid chamber overnight at 4°C. The next day, slides were thrice washed in 1X PBS in a Coplin jar for five minutes each, and then incubated for 1 hour in a humid chamber at room temperature with secondary antibody (FITC anti-mouse, Jackson ImmunoResearch) in the same way as the primary. The slides were thrice washed for 5 minutes in 1X PBS and allowed to dry in the dark at room temperature. The slides were mounted in Vectashield supplemented with DAPI with a glass coverslip. Chromosome spreads were visualized on a Zeiss Axioskop II mot-plus microscope.

### Developmental assays

Genetic isolation of the Y chromosome was performed as previously described (Paredes and Maggert 2009a). Fifteen to twenty male and female adults were placed in a plastic collection bottle (a plastic fly bottle punctured with a 21-gauge needle to provide air holes, and capped with a 35 mm petri plate containing 1% agar made with apple juice from frozen concentrate (Treetop). Females were allowed to lay eggs for 2 days then the plate was replaced with a fresh plate. Around 50 embryos were collected and transferred to 35 mm petri plates containing cornmeal molasses agar, covered, and allowed to develop at room temperature. The plates were analyzed over time, and the developmental stage of the larvae was noted every day for 5 days; this was repeated with several replicate bottles and replicate plates. The same collection procedure was used to collect 1^st^ instar larvae for genomic DNA isolation and Real-Time PCR.

### *18S* stability assay

The experimental design is outlined in (Figure 3). For each independent replicate, 10-20 sibling males and females were allowed to mate for the indicated time periods in vials at room temperature. Once F1 progeny reached the 3^rd^ instar crawling stage for any vial, the F0 adults were transferred to new vials; this was repeated until the desired time period was reached. At the end of each 1, 15, and 30 day period, F0 males were isolated and crossed to virgin C(1)DX females. The adult F1 progeny with the genotype of interest were collected and genomic DNA was isolated from 10-20 flies. As above, 10 ng was used for rDNA quantification.

**Figure 2.**
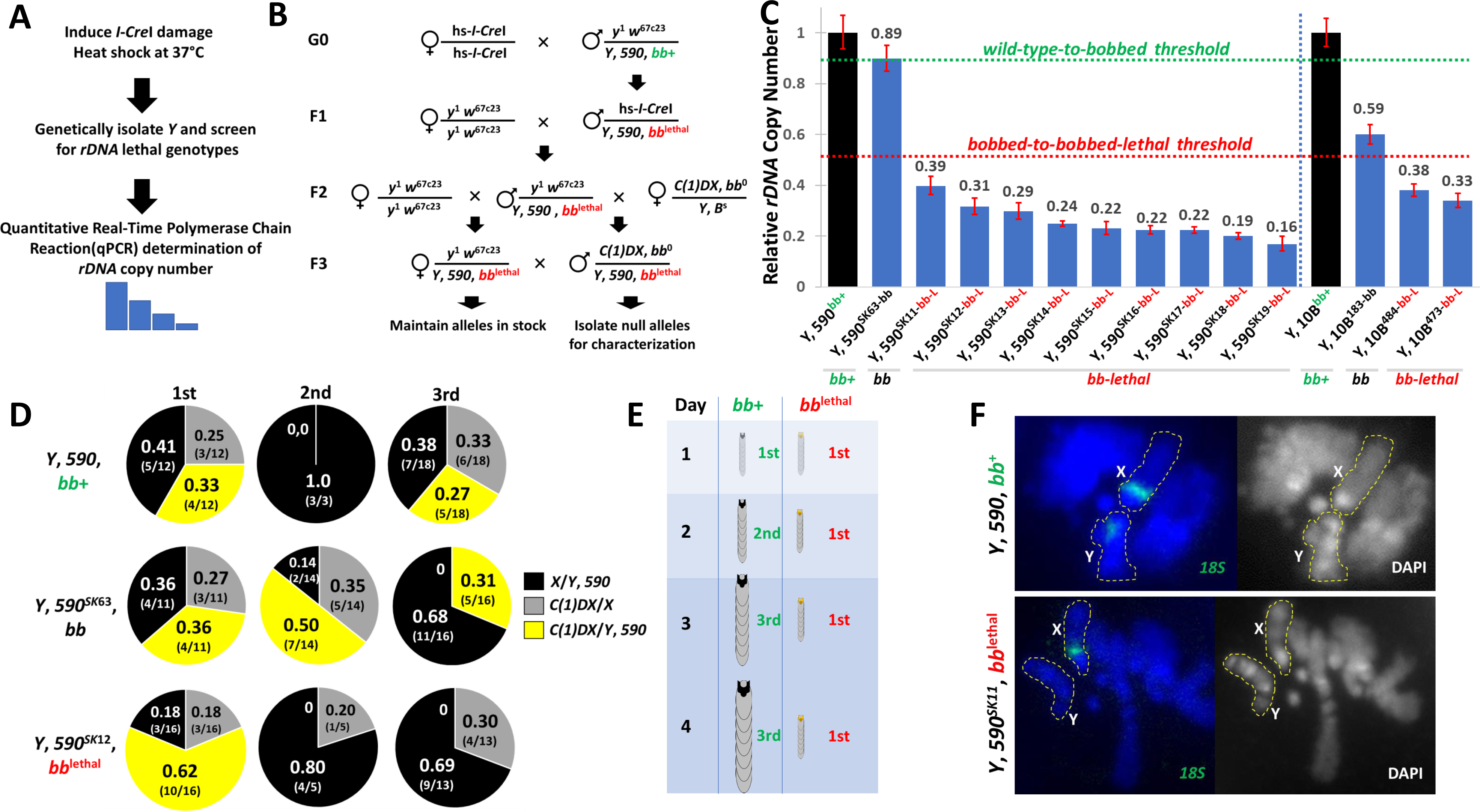
Generation and analysis of *rDNA* deletion alleles. (A) Heat shock-induced expression of I-CreI meganuclease was used to produce DNA damage at the rDNA. Genetic crosses were used to genetically isolate each Y chromosome, which were screened for lethality. Subsequent Real-Time qPCR was used to quantify rDNA copy number. (B) Genetic crosses used to isolate and assess rDNA candidate null alleles. Y, 590 males containing full-length rDNA arrays (bb^+^) were crossed to females carrying an I-CreI construct expressed by heat shock (hs-I-CreI). In F1, male progeny from G0 were crossed to y w females to establish individual lines from different Y chromosome-bearing sperm. In F2, males from established strains were crossed to y w to maintain the strain and to C(1)DX females, which possess no rDNA, to test for lethal or bobbed phenotypes indicative of rDNA deletions. In F3, the individual strains were maintained, and larvae of genotype DX/Y, 590^derivative^ were subject to qPCR rDNA copy number determination. (C) Quantification of the progenitor Y, 590, and deletion derivatives made in this study, and the Y, 10B progenitor and its deletion derivatives from (Paredes and Maggert 2009); copy numbers are relative to tRNA^M-AUG^. These chromosomes help define the wild-type-to-bobbed and bobbed-to-bobbed-lethal copy number thresholds. Error bars are standard deviation. (D) Lethal-phase analysis of rDNA deletions in this study (see also Supplemental Figure 3). Genotypes of individuals were C(1)DX/Y, 590^derivative^. Three different genotypes are shown in the horizontal orientation: Y, 590^bb+^ (wild-type contol), Y, 590^SK63-bb^ (bobbed allele), and Y, 590^SK12-bb-L^ (bobbed-lethal allele). The distribution of the expected sibling genotypes (black, gray, and yellow frequencies) are shown in the vertical orientation as 1^st^, 2^nd^, and 3^rd^ instar larvae. Summary to the right (E) indicates that bobbed-lethal chromosomes (e.g., Y, 590^SK12-bb-L^) terminally-arrest at 1^st^ instar. (F) Cytology of Y, 590 and derivative Y, 590^SK11-bb-L^ reveal no gross aberrations or rearrangements, and that the rDNA deletion chromosomes no longer has rDNA detectable by fluorescence in situ hybridization to the 18S rDNA.

**Figure 3.**
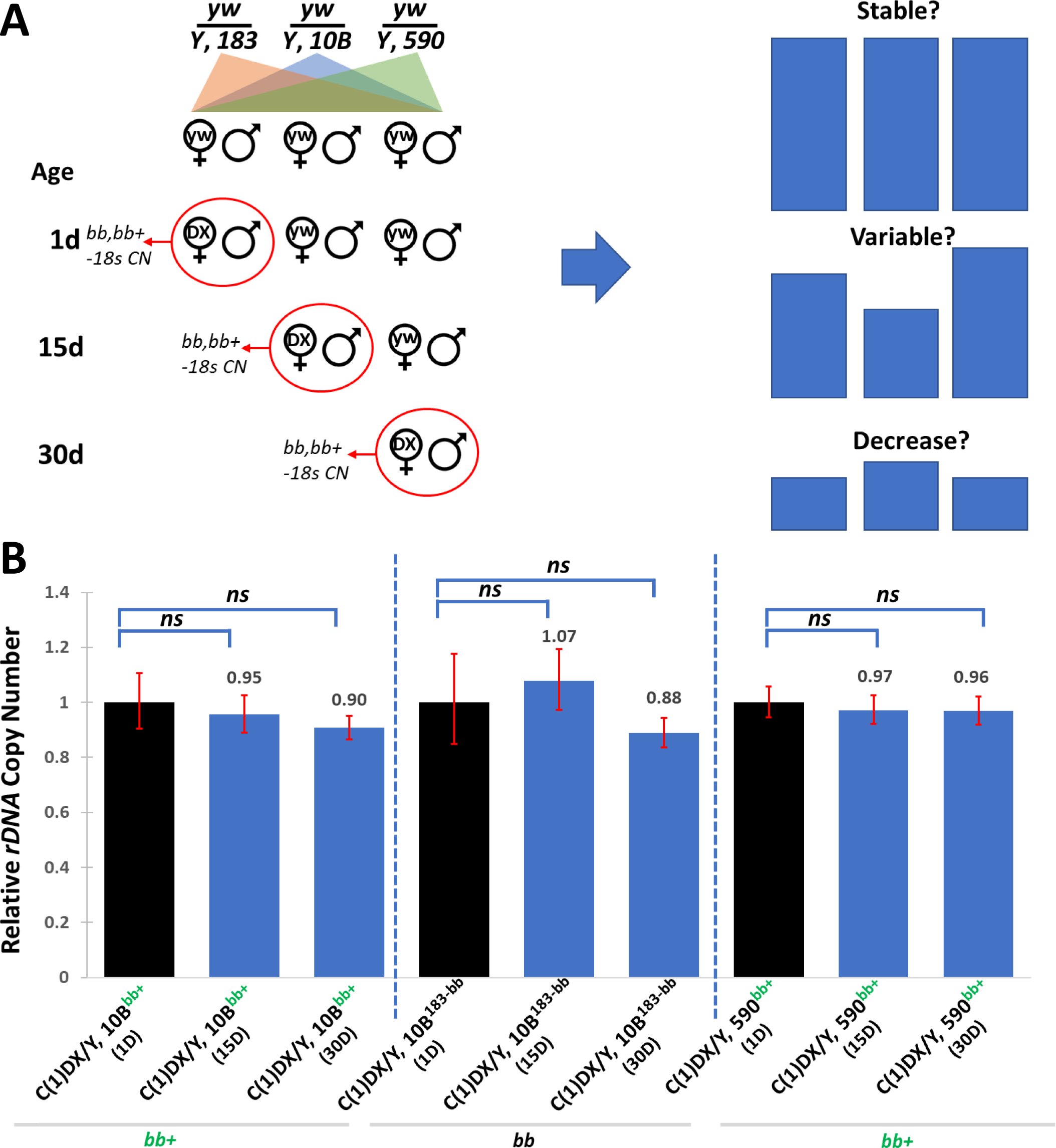
Brood analysis of *rDNA* stability from aging males. (A) Scheme to test sperm from 1-, 15-, and 30-day old males (1 d, 15 d, and 30 d, respectively). Males were co-cultured with sisters until indicated age, then crossed to C(1)DX females, and rDNA copy number assessed in the soma of the offspring. We envisioned three possible outcomes (right): stable rDNA copy numbers from progressively-older sires, increased variability between individuals but relatively-similar rDNA copy number, or loss of rDNA copy number. (B) Three chromosomes (Y, 10B, Y, 183, and Y, 590; Y, 10B and Y, 590 are bobbed^+^, and Y, 183 is bobbed) were brooded for 1 d, 15 d, and 30 d. rDNA copy numbers were calculated relative to the progeny from the 1-day old males (black bars set to 1.0). Statistical test was Student’s t-test, all results showed failure to reject the null hypothesis of equal rDNA copy number; error bars are standard deviation.

### Photography

For body part photography, adults were selected, disassembled, immersed in mineral oil, and illuminated using a Super Bright 10 Klm 12-LED flashlight (iVict). Photographs were taken using a Sony Alpha-III camera attached to a Nikon SMZ 1500 stereoscope. Images were cropped and processed for bright/contrast using Apple Pages.

## Results

To understand the requirements for ribosomal DNA (rDNA), the consequences of retrotranspositional insertion of the rDNA, and the conditions under which rDNA magnify, we first created otherwise-isogenic Y chromosomes that differed only in the extent to which they were deficient for some or all of their 35S rDNA. We began with a single male bearing Y, P{y^+mDint2^ w^BR.E.BR^=SUPor-P}KV00590 (henceforth, “Y, 590”, or Y, 590^bb+^), a Y chromosome possessing a single P element insertion in cytological band h17 in the proximal pericentric heterochromatin of the long arm (Konev et al., 2003).

Apart from the presence of the P element, Y, 590 is a natural chromosome with no rearrangements or polymorphisms detectable by cytology (Figure 1B). This distinguishes Y, 590 and its derivatives from those previously analyzed for rDNA behavior (Y, 10B and its derivative Y, 10C), which had supernumerary rDNA and duplications of X sequence (Paredes and Maggert 2009a; Maggert and Golic 2005). Single Y, 590 chromosomes were sufficient to confer fertility to males, and viability and fertility to females of genotype C(1)DX, y^1^ f^1^ rDNA^0^/Y, 590 (C(1)DX, y^1^ f^1^ rDNA^0^ is henceforth, “C(1)DX”), indicating that Y, 590 is free from any consequential mutations or defects, including at the rDNA.

Due to position effect variegation (PEV) induced by the heterochromatin of the Y, expression of the P-element-containing yellow^+^ and white^+^ genes variegated to an extent that dorsal cuticular cells expressed ∼20 yellow+ spots in X/Y males, and eyes expressed a white-variegating phenotype ranging from ∼10 ommatidia to ∼50% of the eye expressing a white+ phenotype (Figure 1C). The PEV pattern was generally salt-and-pepper, with occasional clonal patches of pigment (Supplementary Figure 1). Due to trans-suppression of PEV by extra Y-linked heterochromatin, aneuploid females (X/X/Y or C(1)DX/Y/Y) and males (X/Y/Y) were readily apparent in our experiments, allowing us to easily monitor chromosome mis-segregation or loss as a result of damage to the rDNA (Bridges and Brehme, 1944; McKee and Karpen, 1990; Maggert, 2014).

### Generation of *de novo* Ribosomal DNA Deletion Alleles

We first analyzed the rDNA array on Y, 590, quantifying 18S rRNA copy number, and the copy numbers of R1-inserted 28S, R2-inserted 28S, and uninserted 28S cistrons. Real-time “quantitative” PCR (qPCR) was used to determine crossing threshold using primers specific for 18S sequence (Figure 1A, primers 8 and 9), and the kinetics of amplification were compared to amplification of the Methionyl-tRNA genes (tRNA^Met-AUG^), as we have done and validated in the past (Paredes and Maggert, 2009a; Aldrich and Maggert, 2014; Guerrero and Maggert, 2011). Using this approach, we estimate that Y, 590 has approximately 130 copies of the 18S gene. Since the 18S is repeated as a subunit with the 5.8S/2S and 28S rRNA subunits, the copy numbers of all three rRNA genes should be the same. For comparison to our previous work (Paredes and Maggert, 2009a, b; Aldrich and Maggert, 2015) and to the known variation in rDNA copy number within natural and laboratory strains (Lyckegaard and Clark, 1991; Long and Dawid, 1980; Stage and Eickbush, 2007; Greil and Ahmad, 2012; Lu et al., 2018; Zhou et al., 2012) we selected five unrelated laboratory strains and determined their rDNA and R1/R2 copy numbers using primers specific for uninserted, R1-, and R2-inserted 28S rRNA genes (Figure 1A, primers 1-7). Analysis showed that Y, 590 is within the range of variation of the rDNA (Figure 1D), consistent with our determination that when Y, 590 is the sole source of rDNA, the organisms do not express a bobbed phenotype, and the rDNA copy number of Y, 590 is above the previously-defined bobbed-lethal and bobbed-viable thresholds (Paredes and Maggert, 2009a).

To create Y, 590 derivative chromosomes that had most or all of their rDNA deleted, we crossed females bearing a heat-shock-inducible I-CreI homing endonuclease to males of genotype y^1^ w^67c23^/Y, 590 (Figure 2A-B) (y^1^ w^67c23^ is henceforth “y w”). Offspring were heat-shocked according to various regimens, and surviving offspring (I-CreI/y w females and I-CreI/Y, 590 males) assessed for male and female survival (Supplemental Figure 2); it has been previously shown that females are less-sensitive to somatic rDNA damage from I-CreI than are males, allowing us to use the male:female ratio or the presence of bobbed males as a proxy for efficacy of I-CreI expression and the amount of rDNA-linked DNA damage (Maggert and Golic, 2005; Paredes and Maggert, 2009a).

There was a clear relationship between heat shock time (I-CreI expression) and male survivability. Three regimens produced few males, but they were sterile (regimens 6, 7, and 8, in red). Instead, we proceeded with regimen #5 (in yellow), outcrossing the males to y w females, and establishing individual lines from each sperm (Figure 2B). Using this approach, we recovered ten new rDNA deletions (generically referred to as Y, 590^derivative^). One (Y, 590^SK63-bb^) expressed a severe bobbed phenotype (as C(1)DX/Y, 590^SK63-bb^ females), and nine expressed a bobbed-lethal phenotype (as C(1)DX/Y, 590^rDNA-^ bobbed-lethal females).

Once established, males from each line were crossed to y w to maintain the strain, and to C(1)DX/Y, B^S^ to quantify rDNA loss using qPCR. C(1)DX/Y, 590^rDNA-bobbed-lethal^ female larvae were collected and total genomic DNA was extracted. qPCR analysis confirmed loss of rDNA copy number on the Y chromosomes of each strain, which was consistent with rDNA loss in previously-published strains from a different parental Y chromosome in terms of copy number and bobbed/bobbed-lethal phenotypes (Figure 2C).

### Characterization of *de novo rDNA* Alleles

After confirming that we had genetically isolated chromosomes bearing candidate lethal rDNA alleles, we sought to identify the developmental stage at which they died from lack of rDNA. We outcrossed +/Y, 590-derivative males to C(1)DX/Y, B^S^ females: this cross produced (i) lethal YY embryos (Y, B^S^/Y, 590^derivative^), (ii) normal XY males (Y, B^S^/+), (iii) triplo-X females (C(1)DX/+), and (iv) females of genotype C(1)DX/Y, 590^derivative^ that possessed the Y as the sole source of rDNA. Owing to the extreme variegated yellow^+^ of the P element of Y, 590, C(1)DX/Y, 590^derivative^ and C(1)DX/+ triplo-X females could be distinguished by the yellow+ phenotype of the latter’s mouth hooks. Larval stages were identified by assessing the appearance of anterior spiracles: absent spiracles were taken as 1^st^ instar, clubbed spiracles as 2^nd^ instar, branched spiracles as early 3^rd^ instar, and everted spiracles as late 3^rd^ instar, respectively. We observed plentiful male and triplo-X female larvae which could be identified at 1^st^ instar, and were seen to continue to develop normally into 3^rd^ instar. However, the C(1)DX/Y, 590^derivative^ female larvae emerged as early 1^st^ instars and remained developmentally arrested at this stage for several days before finally dying (Figure 2D, 2E, Supplementary Figure 3).

The growth arrest of the lethal lines as 1^st^ instar larvae precluded our ability to further investigate the rDNA deficient alleles genetically or cytologically. We therefore assessed the structure and morphology of the lethal rDNA-deficiencies in males of genotype y w/Y, 590^derivative^. Upon inspection, the Nucleolar Organizing Region (NOR) – the cytological locus of the rDNA – of these chromosomes was unremarkable. We attempted to quantify rDNA copy number by length measurements of the NOR secondary constriction of the Y, but could detect no statistically significant correlation between the constriction and qPCR quantification (Supplemental Figure 4A, 4B), leading us to conclude that physical measurement of the secondary constriction is an insensitive and unreliable method of rDNA quantification.

We also attempted to measure rDNA copy number by integrated fluorescence of the rDNA using DNA fluorescence in situ hybridization with probes specific to the 35S rDNA. This approach has been used by others to monitor rDNA dynamics in single cells (Yadlapalli and Yamashita, 2013). We observed signal mostly on the X-linked rDNA compared to the Y-linked rDNA in y w/Y, 590^derivative^ males, and no detectable signal on the Y-linked rDNA of the lethal lines (Figure 2F). This is in contrast to our qPCR data, which indicated those chromosomes possess about 20% of the rDNA as does Y, 590. This discrepancy is likely due to non-linear and/or non-zero thresholds for detection using fluorescence for quantification. Although we could detect population-level differences in average fluorescence in strains with rDNA loss, the variation of fluorescence within an individual vastly exceeded differences between individuals with different rDNA copy numbers (Supplemental Figure 4C, 4D). We therefore conclude that fluorescence quantification is an insensitive and unreliable method of rDNA quantification.

We confirmed that there was not unanticipated contribution to rDNA from the C(1)DX chromosome by collecting yellow female progeny from a cross between C(1)DX/Y, B^S^ females and X^Y/0 males. The C(1)DX/0 progeny also died as first instar, and qPCR of the dying larvae had no detectable rDNA (Supplemental Figure 5).

### Stability of *rDNA* Alleles

Others have reported an intrinsic instability of rDNA copy number on both X and Y chromosomes, resulting in what Lu and colleagues (Lu et al., 2018) likened to the “magnification” characterized by Ritossa (Ritossa, 1973), and Hawley and Tartof (Hawley and Tartof 1985), although it remains unknown if the phenomena are mechanistically related (Paredes and Maggert, 2009a). Our laboratory has produced a number of rDNA deletions – some first published in 2009, some from this study – and multiple analyses over the course of months or years has failed to demonstrate any detectable loss or gain in rDNA copy number by enhancement or suppression of the bobbed phenotypes or by direct qPCR-based rDNA quantification (Aldrich and Maggert, 2014; Aldrich and Maggert, 2015). We therefore sought to directly test rDNA copy number recovery under the same conditions in which it has been described by Lu and colleagues.

We crossed C(1)DX/Y, B^S^ females to males bearing one of three Y chromosomes: Y, 10B (a Y chromosome bearing a full-length rDNA array, also referred to as Y, 10B^bb+^), Y, 183 (a Y chromosome with a severe but viable bobbed allele, derived from Y, 10B, also referred to as Y, 10B^183-bb^), and Y, 590. We did a brood analysis, sampling sperm from 1 day old males, 15 day old males, and 30 day old males (Figure 3A). We crossed individual males to y w virgin females, and collected male offspring 1-2 days after eclosion and divided them into three groups. The first group were mated to C(1)DX/Y, B virgins for 5 days. At the end of those 5 days, the adults were discarded and the progeny allowed to develop to adulthood. Female offspring (C(1)DX/Y^”young”^) were processed for rDNA quantification. The second group of males were kept housed with their sisters for 15 days, allowing them to mate ad libidum to continuously diminish stored sperm. After 15 days, the males were transferred to fresh vials with C(1)DX/Y, B virgins. This second group of males mated for five days, then all adults were discarded, and again female offspring (C(1)DX/Y^”middle-aged”^) offspring were processed for rDNA quantification. The third group were kept housed with their sisters for 30 days, then transferred to new vials with C(1)DX/Y, B, allowed to mate for five days, discarded, and the female offspring (C(1)DX/Y^”old”^) processed. We expected one of three outcomes (Figure 3A): stable rDNA would result in females with rDNA copy numbers equal to the parental Y chromosomes, variable rDNA would result in increased variance with little or no overall loss, and decreased rDNA would appear as detectable reduction of the average. Lu and colleagues reported a 25% loss after by 20 days. We detected no statistically significant decrease in rDNA copy number from any chromosome after any time point (Figure 3B).

We also monitored the enhancement or suppression of bobbed phenotypes, which should occur if rDNA copy number is dynamic in the germline. Although we see variability in the bobbed phenotype of Y, 590, most likely due to its comparably high retrotransposon load, we could detect no obvious trend in either enhancement or suppression of the frequency of bobbed offspring in the other Y chromosomes. As shown in Supplemental Figure 6, there was high variability in the number of bobbed flies, but increases were offset by decreases in the number of bobbed, consistent with the well-known variable expressivity of this phenotype. These results are consistent with our previous work, which showed no changes in rDNA copy number apart from a small increase one or two generations after rDNA damage induction by I-CreI exposure. Stability of the rDNA from one generation to the next appears the norm.

### Analysis of *R1* and *R2 rDNA*-Resident Retrotransposable Element Copy Number

R1 and R2 copy numbers and transcriptional dynamics have been studied extensively in natural and laboratory populations, yet little is known about R1 and R2 in rDNA deficiency alleles (Terracol 1987). We quantified R1 and R2 copy numbers by taking advantage of the specific insertion sites for R1 and R2, irrespective of common 5’ truncations (Figure 1A).

We quantified total, uninserted, R1-inserted, and R2-inserted rDNA copy numbers from our Y, 590-derived bobbed and bobbed-lethal lines (Figure 4A-B). Total rDNA were reduced, consistent with the appearance and severity of the bobbed-lethal phenotypes when these chromosomes were made sole source of rDNA. In contrast, the copy numbers of R1 and R2 were relatively unchanged in deletion alleles, even in extreme cases. The reason for this is unknown, but we consider two possibilities. First, that retrotransposon-inserted rDNA are separate from the uninserted rDNA, including that the C(1)DX chromosome may bear a separate array of R1 and R2 insertions. Second, that I-CreI has a preference for uninserted rDNA cistrons, however it is not clear why that should be the case. It is known that inserted rDNA cistrons are expressed (Eickbush et al., 2008), although perhaps not to the same level as uninserted cistrons (Guerrero and Maggert, 2011). Further, the insertion sites for R1 and R2 are some 840 base pairs from the I-CreI cleavage site, and insertion of retrotransposons does not affect the core or extended I-CreI consensus. We imagine that I-CreI cleaves the rDNA sequence multiple times in each rDNA array, so even inserted rDNA cistrons should be lost as interstitial deficiencies during repair. Nonetheless, our results provide clear proof that the bobbed phenotype is affected by uninserted, but not inserted, rDNA cistrons. Note that we could not discriminate between singly-inserted (that is, containing R1 or R2) and doubly-inserted (containing both R1 and R2) cistrons, however others ‘work has indicated that such double-inserts are relatively rare (Ye et al., 2005).

**Figure 4.**
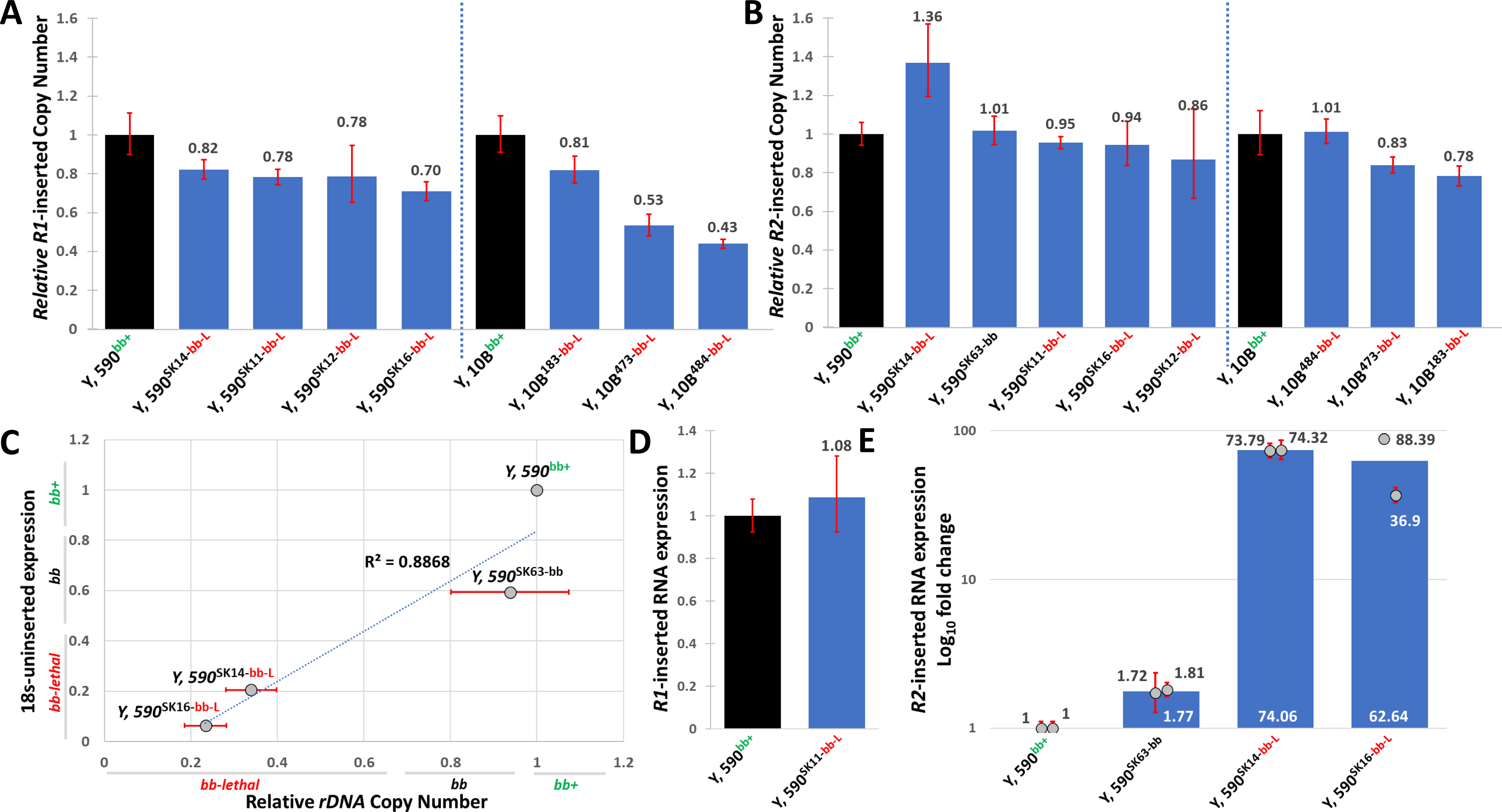
Analysis of *R1* and *R2* copy numbers in *rDNA* deletion alleles. (A) Copy numbers of R1 retrotransposable elements in Y chromosome-linked rDNA arrays from this and a previous study (Paredes and Maggert 2009a) show relative stability of R1 copy number on Y, 590 and its deletion derivatives. Copy numbers are relative to tRNA^M-AUG^ copy number and are reported as fraction of the progenitor Y, 590 or Y, 10B chromosomes. Error bars are standard deviation. (B) As in (A), but showing data for the R2 retrotransposable element copy numbers. (C) Expression of 18S, here determined by steady-state rRNA quantification, is a well-correlated (R^2^ = 0.8868) function of rDNA copy number over a wide range of rDNA copy numbers in y w/Y, 590^derivative^ males. (D) R1 expression in wild-type (y w/Y, 590) and a severe bobbed-lethal rDNA allele (y w/Y, 590^SK11-bb-L^) alleles. (E) R2 expression in wild-type (y w/Y, 590), a mild bobbed rDNA allele (y w/Y, 590^SK63-bb-L^), and two severe bobbed-lethal alleles (y w/Y, 590^SK14-bb-L^ and y w/Y, 590^SK16-bb-L^) alleles.

We quantified 18S expression by reverse-transcriptase qPCR, and found that its expression was greatly decreased in deletions of the rDNA, regardless of the relative stability of R1 and R2 copy number. In fact, expression was directly proportional to 18S copy number (Figure 4C). Thus despite R1 or R2 in the rDNA, their presence does not strongly affect expression of the rRNAs from the uninserted copies within the rDNA arrays.

### Regulation of *R1* and *R2* Retrotransposable Elements

To understand how R1 expression is influenced by the rDNA array, we quantified expression in our rDNA deletion alleles. Neither absolute expression of R1 mRNA nor expression-per-copy of rDNA-resident R1 retrotransposable elements were significantly altered (Figure 4D), suggesting that R1 expression is autonomous and effectively repressed in all conditions.

In contrast, R2 expression was dramatically increased in the rDNA deletions (Figure 4E). In the relatively small deletion expressing a bobbed phenotype, we observed a 1.77-fold increase in steady-state R2 mRNA levels; this is despite any co- or post-transcriptional degradation of R2 (Eickbush and Eickbush, 2014). In two independent bobbed-lethal strains (C(1)DX/Y, 590, rDNA^SK14-bb-L^ and C(1)DX/Y, 590, rDNA^SK16-bb-L^), expression was increased about 70-fold. Previous work by Eickbush and colleagues comparing expression between different Y-linked rDNA arrays identified a correlation between the clustering of the R2-containing rDNA and total R2 expression (Eickbush et al., 2008). In that work, the reason for the differences could not be assertively determined as the Y chromosomes were from different strains. In our study, we can ascribe derepression of R2 directly to the deletion of uninserted rDNA copies because the increased expression was a consequence of rDNA loss from a common single progenitor chromosome. We interpret these data to mean that the R2 elements, in contrast to R1, are regulated at least in part by a global mechanism that is sensitive to uninserted rRNA expression. It is known that R2 elements are repressed by SUMO (Luo et al., 2020), CTCF (Guerrero and Maggert, 2011), and the RNA Polymerase I initiation complex (Fefelova et al., 2022), but it seems likely that those mechanisms of repression are downstream of whatever roles R2 clustering or proximity to uninserted rDNA expression may play (Zhou and Eickbush, 2009).

### Analysis of the Structure of the *rDNA* Loci

To understand the arrangement of R1- and R2-inserted rDNA in the X- and Y-linked arrays, we analyzed their localization using fluorescence in situ hybridization (FISH). Probes directed to the 18S hybridized only to the X-linked array on our bobbed-lethal Y, 590^derivative^ chromosomes (Figure 2F), indicating that the lower limit of detection for FISH is the approximately 20 copies that remain in those strains. In contrast, both the R1 and R2 probes localized to the X- and Y-linked rDNA arrays, whether in wild-type or bobbed-lethal strains. Comparing the wild-type to the bobbed strains, the appearances of R1 and R2 patterns were identical.

The appearance of hybridization allowed us to infer the relative arrangement of the 18S, R1-, and R2-inserted elements within the arrays. The R1 signal localized mainly to the distal X-rDNA– heterochromatin boundary on the long arm, indicating that within the resolution of FISH, the R1 elements are clustered into a single locus (Figure 5A). In incompletely-condensed (i.e., early prometaphase) chromosome spreads, two R2 FISH signals were evident flanking the rDNA on both the X and Y chromosomes (Figure 5B and 5C, respectively).

**Figure 5.**
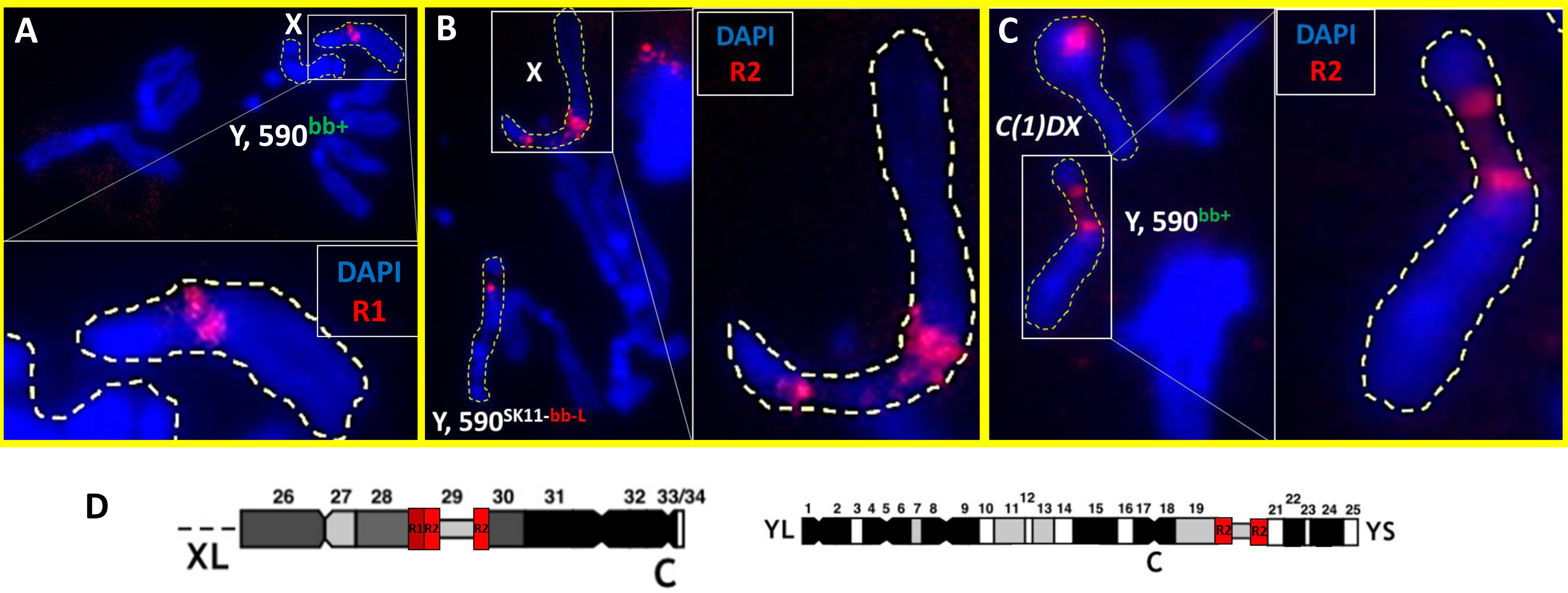
Structural analyses of *rDNA* loci using fluorescence *in situ* hybridization. (A) Mitotic neuroblast spreads highlighting the X chromosome in y w/Y, 590, with FISH signal directed at R1 retrotransposable elements (red). (B) As (A), but FISH signal directed at R2 retrotransposable elements in y w/Y, 590^SK11-bb-L^ males whose deleted Y-linked rDNA forces expression of the X-linked rDNA as indicated by the pronounced secondary construction. (C) As (B), but FISH signal directed at R2 retrotransposable elements in C(1)DX/Y, 590^bb+^ females whose deleted X-linked rDNA forces expression of the Y-linked rDNA. (D) Summary of the results from (A)-(C), plus other spreads (not shown). The X- and Y-linked rDNA arrays (secondary constrictions) are flanked by R2-rich sequence. Further, the X chromosome array has a single locus enriched for R1 sequence that lies distally to the distal R2-rich locus.

We found that the distal Y-linked R2 array was dimmer in fluorescence than was the proximal array. In a few chromosome spreads, the sister Y chromatids were adequately separated so as to be independently identifiable. In those cases, it was unusual for the two sister rDNAs to have the same level of fluorescence, underscoring the insensitivity of FISH for quantification. Since these arrays should be identical products of replication, we interpret the difference to be due to differential probe hybridization, chromatin structure, or some other factor. However, relative fluorescence between proximal and distal foci was consistent, thus we believe that the distal R2 clusters likely contain fewer copies than do the proximal R2 clusters. Alternatively, it is possible that the distal R2 clusters have a lower degree of homology to the probe, or the chromatin structure at the distal R2 cluster is less accessible to in situ hybridization.

The arrangement we see is in good agreement with Eickbush’s “domain” model of retrotransposon regulation, in which he posits that most of the R1 and R2 inserted copies are packaged into the heterochromatic regions flanking the active uninserted rDNA (Eickbush et al., 2008) (Figure 5D), contributing to their silencing. Our results are also similar to the situation described for the Chinese silkworm Bombyx mori, where R1 and R2 signals localized to the heterochromatic flanks of the rDNA arrays (Pérez-González and Eickbush, 2001).

We were surprised to detect intense R1 and R2 signals on the C(1)DX chromosome (Supplemental Figure 7), which has been reported (and we confirmed) to be devoid of any active rDNA (Supplemental Figure 5). Normally, these two retrotransposons are thought to be obligate residents of the rDNA, however no rDNA were detectable using FISH or qPCR, the C(1)DX chromosome was never seen to exhibit a secondary constriction, and C(1)DX/0 flies are lethal very early in development owing to rDNA loss. We therefore conclude that the C(1)DX chromosome contains an island of R1- and R2 retrotransposon sequences outside of the context of the rDNA. It is not clear if this is an artifact of the production of the C(1)DX compound chromosome, is a vestige of the original X chromosomes from which C(1)DX was generated, or if even wild-type chromosomes possess “free” R1 and R2 elements outside of (but perhaps adjacent to) the rDNA arrays.

### Ribosomal DNA Dominance: Secondary Constriction and *Histone H3.3* Incorporation

In most metaphase and anaphase spreads of neuroblasts taken from third instar larvae bearing wild-type Y, 590 chromosomes, the fluorescence signals from the X-linked R2 were very close together. Although they were discriminable, they had little or no sign of a constriction between them. We interpreted this to mean a high degree of condensation of the uninserted rDNA, revealing a generally inactive X-linked rDNA array. However, in some chromosome spreads it was apparent that the two X- linked R2 clusters were widely separated. In those cases, the rDNA had a classic constriction, meaning a reduced width and staining, and an extended length, at the rDNA locus. The appearance of a constriction is accepted to mean under-condensed chromatin as a result of recent or ongoing transcription of rDNA cistrons. To confirm this interpretation, we employed a histone H3.3-GFP fusion transgene (Ahmad and Henikoff, 2002; Greil and Ahmad, 2012) to preferentially label rDNA arrays in prometaphase and metaphase that had been expressed in the the previous G1-phase. Despite efforts, we were unable to detect Histone H3.3-GFP fluorescence in living cells, so instead performed immunofluorescence to the GFP moiety. Fluorescence decorated the rDNA constrictions and was concomitant with the separation of R2-containing clusters flanking the X-linked actively-expressed rDNA.

We interrogated nucleolar dominance relationships in our bobbed-lethal lines by assessing the degree of secondary constriction formation and Histone H3.3-GFP incorporation in y w/Y, 590^derivative^ males. We scored each mitotic figure for the level of fluorescence staining on the X- and Y-linked rDNA arrays, sorted into one of five categories (Supplemental Figure 8) depending on whether the expression was exclusive to one chromosome, heavily biased to one array, or comparable between the two. We have already determined that FISH intensity is an insensitive means of measuring rDNA copy number, and under that same logic, we chose a conservative interpretation of Histone H3.3-GFP incorporation (green spots) by grouping exclusive and dominant expression into the same category. In support of this decision, we saw positive signal on the X-linked rDNA in most spreads and observed an overall spectrum of dominance relationships between the X and Y arrays (Figure 6A, C).

**Figure 6.**
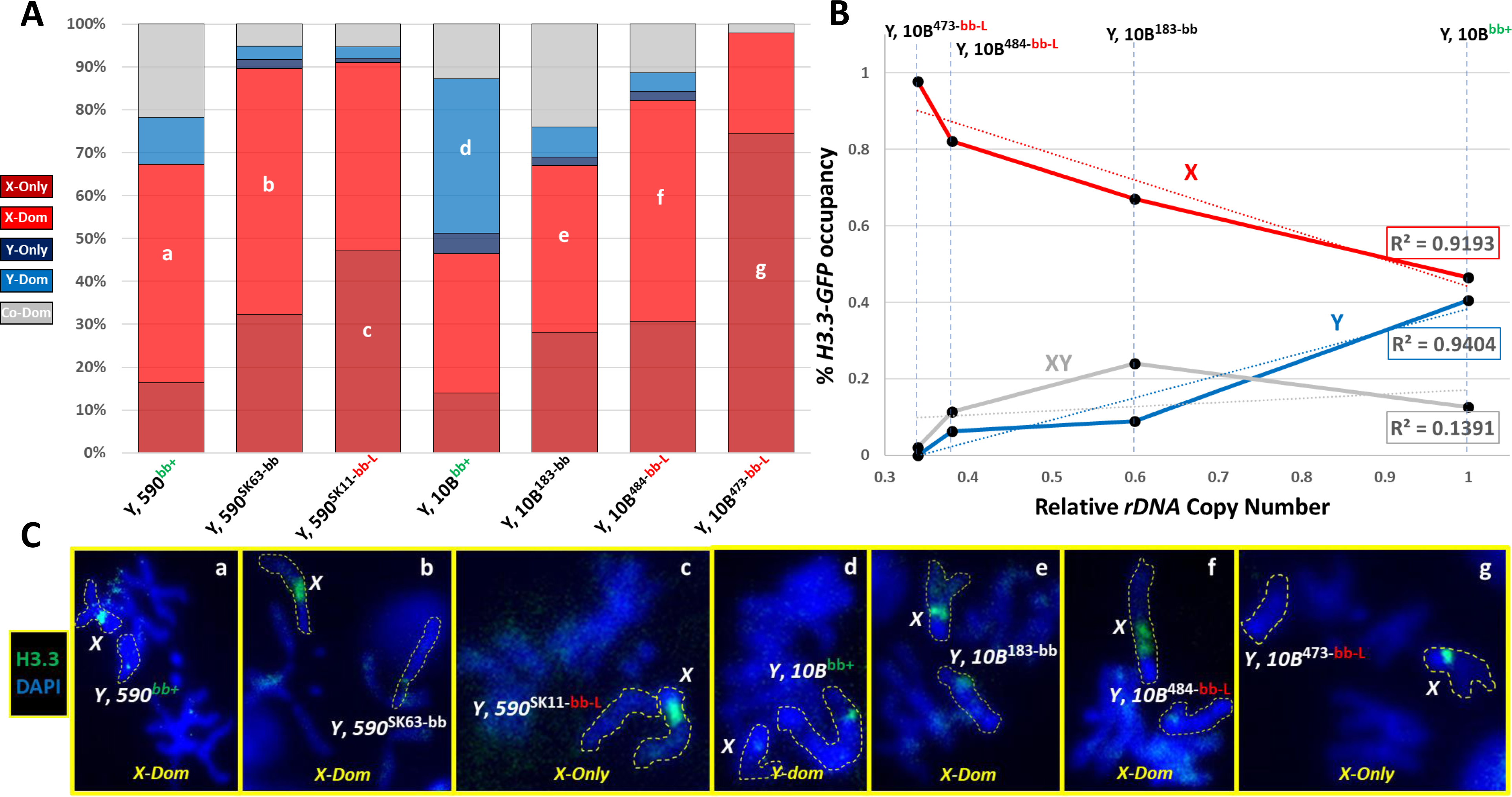
*Histone H3.3* incorporation at the *rDNA* loci of *rDNA* deletion alleles. (A) Distribution of nucleolar dominance states in wild-type and derivative bobbed Y chromosomes. Each mitotic figure was independently assigned to one of five categories, depending on the relative intensity of Histone H3.3 incorporation (in green) at the rDNA – X- and Y-exclusivity, X- and Y-bias, and co-dominance (see Supplemental Figure 8). Letters refer to representative images shown in (C). (B) Correlation between rDNA copy number (X-axis) and biased or exclusive Histone H3.3 incorporation as an indication of nucleolar dominance. Expression of the X-linked rDNA and Y-linked rDNA regress well with Y-linked rDNA copy number (red and blue lines, respectively), such that low rDNA copy number on the Y is correlated with X-linked rDNA activity. (C) Representative images of X- and Y-inked rDNA expression; lower-case letters refer to genotypes and dominance categories shown in (A).

The X-linked rDNA arrays in males bearing the Y, 590^rDNA-bobbed-lethal^ chromosomes were consistently constricted and showed incorporation of Histone H3.3-GFP. This indicates constitutive expression of the X-linked array despite the otherwise-normal dominance of Y-linked rDNA. In contrast, males of genotype y w/Y, 590^rDNA-bobbed^ exhibited inconsistent constriction and Histone H3.3-GFP incorporation, indicating that derepressed X-linked arrays responded to the status of expression and rRNA output from the Y-linked array. Specifically, we recorded a positive correlation between rDNA and Histone H3.3-GFP incorporation on the Y, which we interpret as evidence that if the Y-linked rDNA has sufficient rDNA for translational demands, it is the preferred array in males. We also recorded a negative correlation between rDNA copy number on the Y and Histone H3.3-GFP incorporation on the X (Figure 6B), which we interpret as evidence that as the Y-linked rDNA array is less capable of supplying necessary rRNAs, the X linked array becomes active. This was expected due to the genetically recessive nature of both X- and Y-bobbed alleles, however this is the first direct cytological evidence of this switch between Y dominance and X-expression in Y-linked rDNA mutations. Our system of otherwise-isogenic rDNA deletion alleles allows us to conclude that loss of rDNA on the Y is causal for this switch. Moreover, the variation in X versus Y-dominance appeared cell-by-cell within spreads of cells from single brains, indicating that expression of the X-linked array was sporadic and reversible, and was not a permanent derepression caused by structural changes to the X-linked rDNA array, or due to specific genetic interactions between some X-linked and Y-linked rDNA arrays.

Although we saw clear bias and switching in our strains, the activity at the X-linked rDNA array was very common in all our strains. This is in contrast to what was reported previously by Greil and Ahmad (Greil and Ahmad, 2012) and Warsinger-Pepe and colleagues (Warsinger-Pepe et al., 2020), which showed a stronger Y dominance bias. We presume that the reason is due to the structure of the specific X and Y chromosomes used in these three different laboratories. Greil and Ahmad reported some strains with permanently-altered dominance patterns, yet our results here show that rDNA copy number loss can erase or bias dominance. When we analyzed the X and Y bias of the chromosomes used by Greil and Ahmad, we recapitulated their observations (Supplemental Figure 9). While it seems clear that extended derepression of an X-linked array by Y-linked rDNA loss can lead to permanent changes in the chromosome (Paredes and Maggert 2009a, b; Aldrich and Maggert, 2015), we did not detect such a transition in our chromosomes in the relatively short time frame of this study.

### Conditions Affecting *rDNA* Magnification

Earlier studies to differentiate between pre-meiotic and meiotic magnification employed mutations in genes necessary for DNA repair and recombination. It was observed that magnification was prevented in genetic backgrounds with mutations in post replication repair proteins mei41, which encodes a repair kinase, mus101, which encodes a topoisomerase-II binding protein, and mus108, which has not been identified but confers mutagen sensitivity and blocks normal meiotic recombination and repair. These genes are required for DNA repair and for magnification at the rDNA. It is likely that other DNA repair enzymes are required for magnification based on the currently-accepted consensus model, which envisions increased DNA damage at the magnifying rDNA, illicit sister-directed repair, and unequal sister chromatid exchange (Endow et al., 1984; Kindelay and Maggert, 2023).

We proposed elsewhere that magnification in males occurs when the normally-silent X-linked rDNA array is derepressed (Kindelay and Maggert, 2023). This led us to hypothesize here that conditions that disrupt “normal” nucleolar dominance, either by specifically affecting dominance or by generally affecting silencing mechanisms, will lead to increases in magnification. Previous work has focused on mutant conditions that reduce magnification by affecting the requisite mechanisms of repair, while our hypothesis predicts some mutant conditions will induce magnification which, until very-recently, has not been observed.

First, we assessed the ability of our bobbed-lethal alleles to magnify in males in which the dominance relationship had been disrupted. For this, we used the co-dominant X chromosome from Greil and Ahmad’s work – which we call “KX” in this study. We tested a subset of rDNA alleles that included both the bobbed Y, 10B^484-bb-L^ rDNA allele and the bobbed-lethal Y, 10B^473-bb-L^ rDNA allele, as these alleles were expected to give us the best indication of whether or not magnification was occurring even if the increase was small or moderate since these alleles never magnify when made the sole source of rDNA. In both cases, magnification could be identified as suppression or complete reversion of the bobbed phenotype (Figure 7A). We created KX/Y males, using a known co-dominant X chromosome (Greil and Ahmad, 2012), and crossed them to C(1)DX/Y, B^S^ females and screened for living female offspring, which would be the product of rDNA magnification (Figure 7B).

**Figure 7.**
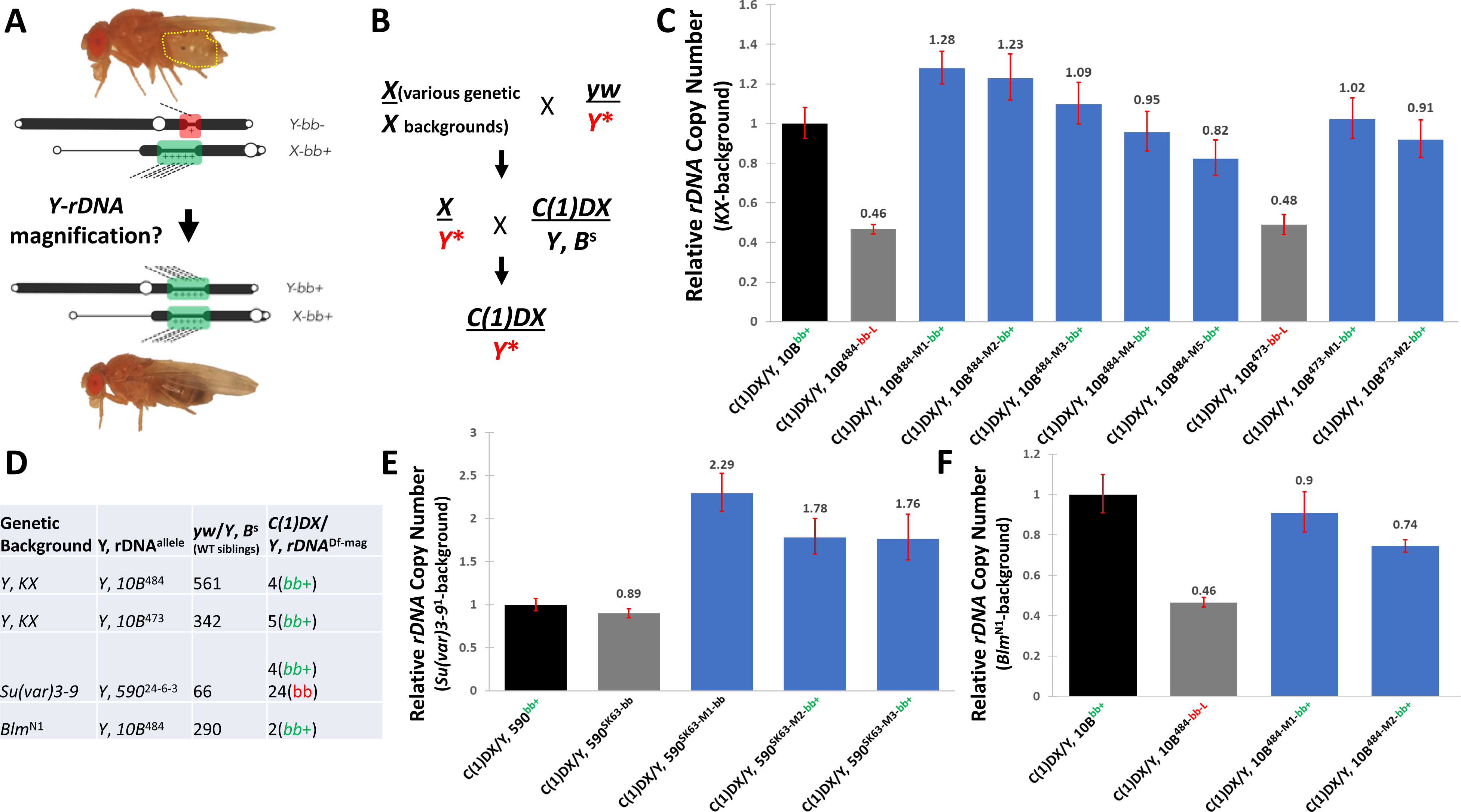
*rDNA* Magnification caused by mutations altering *rDNA* regulation. (A) Adults can express a bobbed phenotype if their rDNA is insufficient for translational demands (top), and magnification to higher rDNA copy number will revert this phenotype to wild-type (bottom). (B) Females of varied genetic constitution were crossed to males with a Y chromosome to be tested for magnification (red Y*). Male progeny with the mutation and Y chromosome were crossed to females of genotype C(1)DX/Y, and female progeny assessed for bobbed+ indicating magnification. (C) The KX chromosome was tested for its ability to magnify the Y, 10B^484-bb-L^ and Y, 10B^473-bb-L^ chromosomes. Black rDNA copy number data show control Y, 10B, the progenitor of Y, 10B^484-bb-L^ and Y, 10B^473-bb-L^; “WT” indicates unmagnified chromosomes that were not placed in the presence of the KX chromosome; “bb” indicates chromosome in progeny that did not exhibit bobbed+; MAG1-5 and MAG1-2 indicate chromosomes isolated from bobbed-revertants. The latter category shows magnification of rDNA copy number. (D) Summary table showing number of revertants from (B), (D) and (E). (E) As in (C), but with the Su(var)3-9 mutation. (F) As in (C) but with the blm/mus309 mutation. Error bars show standard deviation.

We measured rDNA copy number in C(1)DX/Y, 10B^484-bb-L^ controls (whose fathers did not include the KX), female offspring that were severely bobbed and were expected to be escapers, and putative magnified bobbed^+^ females. The first two categories had rDNA copy number consistent with the original Y, 10B^484-bb-L^ chromosome, while the bobbed^+^ chromosomes were all magnified to copy numbers above the bobbed threshold (Figure 7C, D). Subsequent outcross confirmed that these were single magnified Y, rDNA^+^ chromosomes, and not C(1)DX/Y, 10B^484-bb-L^/Y, 10B^484-bb-L^ aneuploids. Notably, we found that some magnified chromosomes contained more than twice the rDNA complement of the unmagnified chromosome – this behavior has been noted before in rDNA magnification (Hawley and Marcus, 1989), and its explanation remains unknown. Similar magnification was seen on the Y, 10B^473-bb-L^ progenitor chromosome. Because the Y, 10B^473-bb-L^ chromosome has never before shown magnification to wild-type, but the KX chromosome caused it to, we conclude that disrupted nucleolar dominance is sufficient to induce “magnifying conditions.” This suggested to us that any mutation that derepresses the silencing of the X should equally induce magnification.

Next, we tested for magnification in flies bearing heterozygous mutations in factors known to regulate chromosome pairing and gene silencing. We crossed Y chromosomes into genetic background containing mutations in genes establishing cohesin-dependent silencing – SMC1 and CTCF. SMC1 is a subunit of the cohesin complex and is a member of the Structural Maintenance of Chromosomes (SMC) group of proteins essential for chromosome stability during replication, and segregation during the subsequent anaphase (Dorsett, 2019). No magnification was observed in male flies with the heterozygous genotype y w/Y, 10B^484-bb-L^; SMC1/+. We also examined heterozygous mutants of CTCF, a chromatin insulator with roles in transcriptional activation and repression, which acts through recruitment of cohesin, and has been shown directly to be involved in silencing the rDNA (Guerrero and Maggert, 2011). We could detect no magnification in y w/Y, 10B^484-bb-L^; CTCF/+ flies, although magnification has been reported previously using a different CTCF allele (Guerrero, 2011). From both these crosses we did observe occasional apparent revertants, but subsequent analysis showed them to be Y aneuploids (C(1)DX/Y/Y females) derived from nondisjunction in male meiosis.

We next tested genes involved in heterochromatin formation and maintenance, specifically the Histone H3 Lysine-9 methyltransferase encoded by the Su(var)3-9 locus, and the H3K9-Methyl binding protein encoded by Su(var)205. These genes have been shown to be necessary for the regulation and integrity of the nucleolus and organization of the rDNA in numerous studies (Peng and Karpen, 2007) as heterozygous loss-of-function lesions lead to reduced rDNA copy number in soma and germline (Paredes and Maggert, 2009b; Peng and Karpen, 2007; Paredes et al., 2011; Greil and Ahmad, 2012; Aldrich and Maggert, 2015). The former has also been established to derepress a silent X-linked rDNA, disrupting nucleolar dominance. Our Y, 590^SK63-bb^ bobbed allele exhibited magnification in Su(var)3-9 heterozygotes, as evidenced by the appearance of fully wild-type revertants of bobbed (Figure 7D, E). Those offspring were outcrossed to confirm they were euploid, and then by isolation of rDNA from DX/Y, 590^SK63-bb-rev^ females. In this quantitative test, we again detected greater-than 2-fold increase in the rDNA array size. We conclude that these are bona fide magnification events, though we are cautious to note that we cannot assertively determine if the mechanism of magnification here is the same as that seen in non-mutant conditions (e.g., Hawley and Tartof 1985). In contrast, despite clear and strong loss of rDNA in Su(var)205 heterozygotes, we could detect no magnification of Y, 590^24-6-3^ in y w/Y, 10B^484-bb-L^; Su(var)205/+ males.

Finally, we investigated mutations in blm, a member of the recQ family of helicases. Mutations in these genes cause Bloom Syndrome in humans, which is a rare autosomal recessive disorder characterized by increased genomic instability and susceptibility to cancer (Kaur et al., 2021) owing to Blm’s requisite role in homology-directed repair and replication of difficult-to-replicate DNA. In Drosophila, deficiencies in blm (historically, the mus309 locus) cause defects in embryogenesis (McVey et al., 2007) and an increase in genome rearrangements involving repetitive sequences in the genome (Garcia et al., 2011) possibly due to defects in replication of these regions (Ruchert et al., 2022). In Y, 10B^484-bb-L^; blm^N1^/+ males, we detected an increased frequency of rDNA magnification. Again, we confirmed euploidy and quantified the extent of rDNA magnification using qPCR (Figure 7D, F).

## Discussion

To better understand the biology of the ribosomal DNA arrays (rDNA, the locus of transcription of the 35S rRNAs), we created de novo deletion alleles that, by themselves, were incapable of suppling sufficient rRNAs for translational demands. When the sole source of rRNA, these alleles confer a bobbed phenotype. The alleles of the Y-linked rDNA array were generated from a single otherwise wild-type chromosome, unrelated to the chromosomes that we and others have used for previous characterizations (Paredes and Maggert, 2009a, b; Watase et al., 2022; Nelson et al., 2023; Maggert, 2014). Nonetheless, our findings were fully consistent with the requirements for rDNA to supply ample bobbed^+^ function, indicating that different Y chromosomes may differ in their rDNA, R1, and R2 copy number, but the supply of rRNAs follows a simple linear relationship with the uninserted rDNA copy number. This appears to be a general property of rDNA as the same correlation has been described for Caenorhabditis elegans and Arabidopsis thaliana (Morton et al., 2023; Lopez et al., 2021). For some organisms, the correlation was between copy number and growth (Delany et al., 1994; Su and Delany, 1998; Schneeberger and Cullis, 1991). We saw no evidence that any rDNA copy is alone better able to provide necessary rRNAs, indicating to us that all rRNA genes are essentially equivalent despite some known polymorphisms in rRNA and regulatory sequences, although careful analysis of transcription output by individual cistrons has never been done despite evidence for regulatory heterogeneity (Grimaldi et al. 1990).

### *rDNA* Requirements in Embryogenesis and Larval Life

The rDNA deletion alleles generated in this study allowed us to determine the phenotype phases of limited translational capacity. In organisms with very few (C(1)DX/Y, 590^derivative^) or no (C(1)DX/0) rDNA, the only translation is by maternally-supplied free or ribosome-bound rRNAs. Surprisingly, we found that such rDNA^null^ Drosophila could complete embryogenesis. However, rDNA^null^ larvae were unable to then initiate or complete their first larval molt, so they died days after hatching as unnaturally- old 1^st^ instar larvae. Death of rDNA^null^ larvae was progressive over the course of our experiments, with underrepresentation of rDNA^null^ 1^st^ instars evident on the first day after hatching, and numbers dwindling for the next 3-4 days of our experiments. We suggest that this variance in lifespan may be due to variance in rRNA loading into different eggs, though this has yet to be tested.

We also generated C(1)DX/Y, 10B^473-bb-L^, which has very little rDNA – about 35% of the amount needed for full viability and fertility (Paredes and Maggert 2009a). Female larvae were able to complete both larval molts, making it to 3^rd^ instar. However, they were smaller than their wild-type siblings and died as 3^rd^ instar unable to initiate metamorphosis.

With our work, we can now define four thresholds of bobbed mutants. We define 100% rDNA as the “wild-type” minimum necessary for full viability and fertility in intact organisms, and full cell- viability, such that we can detect neither developmental delays, truncated/bobbed bristles, or cuticular herniations. Below this amount, we observe herniated cuticles and truncated bristles, defining the “wild-type-to-bobbed” threshold. Fewer rDNA, about 80-85% of the wild-type amount, causes under-representation, extreme cuticular defects (including exposed hypoderm), and death, defining the “bobbed-to-bobbed-lethal” threshold. These two thresholds were known from classical work on the rDNA, were defined by ours and others’ previous work (Tartof, 1973; Terracol, 1987) and were recapitulated in this current study. Still less rDNA, approximately 35%, is required for viability, specifically for metamorphosis at the end of the 3^rd^ larval instar, defining the “metamorphosis-bobbed” threshold. Finally, less than this amount of rDNA causes a developmental arrest at 1^st^ instar, defining the “molt-bobbed” threshold. This last threshold appears to be the lowest threshold affected by rDNA copy number, because complete ablation of rDNA (in a C(1)DX/0 organism) behaves identically. These thresholds are defined in females of genotype C(1)DX/0 or C(1)DX/Y, rDNA^deletion^. It therefore does not appear that the non-rDNA portions of the Y chromosome affect these thresholds. As yet we cannot determine if these threshold are the same in males as in females; we have no indication they would not be.

### Stability of the *rDNA*

We and others have reported conditions in which rDNA copy number is unstable, including under dietary stress and activation of the Insulin/Insulin-like pathway (Aldrich and Maggert, 2015), in mutants affecting heterochromatin-induced gene silencing at the rDNA (Paredes and Maggert, 2009b), and in cultured (Xu et al., 2017), and biopsied (Valori et al., 2020) cancer cells. However, others (Lu et al., 2018) have reported a dramatic instability at the rDNA, which they characterize as premeiotic loss in aging germline stem cells, early zygotic recovery, early germline re-loss, mature germline stem cell re-recovery, then another round of age-dependent loss. In this work, investigators propose this multiphasic loss-and-gain cycle as the reason for the known variation in rDNA copy number in wild populations (Dawid and Long, 1980; Lyckegaard and Clarke, 1991). However, this remarkable level of generation-to-generation instability in wild-type flies in normal conditions was unexpected for three reasons. First, old bobbed alleles, some dating back over 100 years (Bridges and Brehme, 1944) and including those used in current studies, have been remarkably stable in terms of phenotype and rDNA copy number. A corollary to this is the evident stability generation-to-generation in recently identified or created strains. Second, analysis of the rDNA from the 1960s through the 1990s did not report instability or age-effects, despite focusing on rDNA copy number and conditions under which they may change. Stability is evident in extensive molecular characterization of individual rDNA cistrons that had been molecularly tagged with P elements (Bianciardi et al., 2012), or with natural retrotranspositional insertions (Zhou and Eickbush, 2009). Third, our own work has not detected any age-effects at the Y-linked rDNA using multiple Y chromosomes (Aldrich and Maggert, 2015; Susamcı et al., in preparation). In this present study, we directly assessed loss of rDNA in the same conditions under which it has been recently reported (Lu et al., 2018). We could detect no changes in uninserted copy number based on qPCR quantification or the appearance of bobbed offspring.

To reconcile these disparate observations, we suggest that the assays used in studies that report dramatic instability (Lu et al., 2018) – mostly nucleoli that are “deformed” or have “atypical morphology” – are poor proxies for rDNA copy number. To wit, we compared fluorescence intensity based copy number determination with PCR-based ones, and find very little correlation; the variation within the former is simply too high to be a reliable metric. Rather, we expect that the “loss” described by others is due to developmental changes to nucleolar morphology, a function of both developmental stage and age, that affect fluorescence in situ hybridization, chromatin compaction, nucleolar size, or the like. It is formally possible that some strains are susceptible to rapid losses and gains, while others are not, however we find no evidence for this. How Y-specific instability of the rDNA can be reconciled with proposed mechanistic models of loss and gain, or with models of population dynamics, is not clear.

In progressively-aged male flies, we detected losses and gains of the retrotransposons R1 and R2 (Supplementary Figure 10A, 10B, 10C). However, the unstable R1s and R2s appear to be lost from extra-rDNA arrays, and so do not represent loss of bona fide ribosomal RNA sequence that alter translational capacity. Likely the source of the lost R1s and R2s is the blocks of elements flanking the rDNA rather than the rDNA itself, which would explain the observed stability of the rRNA-producing rDNA cistrons. We are investigating the reason for R1 and R2 loss, but analysis is complicated by the extra-rDNA R1 and R2 on the C(1)DX chromosome, whose stability dynamics are yet-unexplored.

### *R1* and *R2* Regulation by the *rDNA*

We used new rDNA deletion alleles in conjunction with alleles generated in a previous study, and found a clear linear relationship between the copy number of rDNA and the steady-state level of rRNA in those organisms. The evidence therefore suggests that rRNAs are produced constantly and uniformly from all uninserted rDNA cistrons. Further, that rRNA expression is not subject to compensatory over-expression when rDNA copy number is reduced, arguing that feedback mechanisms to produce constant rRNA from varied sizes of rDNA arrays do not exist. Rather, robustness in ribosome number is accomplished through relatively high degradation of rRNAs during processing and the extremely-long half-life of the rRNAs packaged into ribosomes. We imagine that all uninserted rDNA copies are constitutive and not subject to alteration of transcriptional output. This is a different situation from yeast, where altered elongation rates can compensate for lower rDNA copy number (Schneider et al., 2007), yet it is consistent with work in C. elegans in which some phenotypes exhibit genetic interactions with the rDNA, suggesting that natural variation of the rDNA impacts global translational capacity (Morton et al., 2023). To our knowledge, in Drosophila, the only phenotype known to genetically interact with the rDNA is heterochromatin-induced gene silencing (position effect variegation) (Paredes and Maggert, 2009b; Zhou et al., 2012; Larson et al., 2012), however catalogued changes in mRNA levels certainly leave open the possibility for yet-undiscovered phenotypic variation mapping to rDNA copy number variation (Bughio and Maggert, 2019).

We also found that the amount of R1 expression is relatively constant between full-length rDNA arrays and very short rDNA arrays, even in extreme deletions of the rDNA where the number of R1 elements does not change much. This fits with our cytological observation that detectable R1 elements are clustered at the distal flank of the rDNA array in a way that make them robust to loss or expression. We interpret this to mean that the R1-inserted rDNA cistrons are all independently repressed by some mechanism that is insensitive to uninserted rDNA copy number or expression, or that the R1 elements are sequestered into a structure that globally represses their expression.

In contrast, expression of the R2 retrotransposons is very sensitive to uninserted rDNA copy number, such that R2 expression increases as rDNA copy number decreases. The reason for this is less clear, since the R2 elements also cluster by FISH analysis, though they flank the rDNA constrictions of the X and Y chromosomes rather than being limited to just one side. Hence, the idea that R2 elements are independently repressed by sequence-specific DNA binding repressors is less tenable than the counter-proposal (made by Eickbush and colleagues (Zhou et al., 2013)) that the R2 elements are regulated by clustering. How the R1s and R2s have evolved different mechanisms of regulation is not clear, yet our analysis of rDNA deletions clearly underscores different mechanisms at play. Nonetheless, we detect that uninserted rDNA, R1-inserted rDNA, and R2-inserted rDNA are all subject to fundamentally different modes of regulation.

### Regulation of Nucleolar Dominance

Nucleolar dominance was discovered in interspecific hybrids, and manifests as the production of rRNA from one of multiple rDNA arrays, while the other rDNA arrays are silent. Greil and Ahmad were the first to show that a dominance-like relationship occurs in wild-type Drosophila melanogaster, where the X-linked array is inactive in males and the entire pool of rRNA derives from the Y-linked array. They found certain chromosome pairs that did not obey this dominance relationship, and instead acted co-dominantly, indicating control of dominance by both cis-acting and trans-acting factors. Later work by Warsinger-Pepe and colleagues probed the developmental control of the phases of dominance (Warsinger-Pepe et al., 2020). Neither study identified the features of dominant or co-dominant chromosomes that enforced or maintained the dominance relationship, and Greil and Ahmad argued against a simple measurement of rDNA copy number as the controlling factor using a series of Y chromosomes of different origins.

We found that our full-length Y chromosomes are dominant or heavily-biased in their expression so that most rRNAs in males derive from the Y-linked rDNA array. However, because our deletion alleles are derived from a single “bottlenecked” Y chromosome, we can definitively conclude that as the rDNA is lost, the bias switches from Y-to X-linked arrays. Hence, in our experiments, the amount of rRNA derived from the X chromosome is controlled by the Y chromosome’s adequacy.

We monitored Histone H3.3-GFP incorporation and the formation of secondary constrictions as indicators of rDNA expression. In X/Y male neuroblasts, most individual immunofluorescence images showed Y-linked expression. In males with shorter rDNA arrays, we observed an increase in the fraction of mitotic spreads with X-linked rDNA expression, although there remained some mitotic chromosome spreads with silent X-linked rDNA arrays. This indicates that the Y is constitutively expressed, even if it is unable to supply sufficient rRNA, and further than the X chromosome vacillates between silenced and expressed, depending on the needs of the cell. In other words, a deleted Y-linked rDNA array causes the X-linked array to turn on more often, but expression is still repressed a subset of the time.

How the X-linked rDNA continually or repeatedly senses and reacts to rRNA concentration in a cell is a major question that arises from our work. But our work also provides a clue: we noted that the strength of the dominance relationship is correlated with the number of R1 and R2 insertions. Loss of the rDNA in the deletion alleles generated from Y, 10B causes a derepression of the X, but the strength of the derepression response was less pronounced in the Y, 590-derived rDNA alleles. This latter chromosome has many more copies of R1 and R2 than does Y, 10B, which we propose makes it less sensitive to loss of dominance. Further, the strongest dominance relationship we observed was exhibited by the Y chromosome first characterized by Greil and Ahmad and, as predicted, we found that this chromosome has a very high load of R1 and R2 retrotransposon insertions.

Taken together, we conclude that uninserted Y-linked rDNA cistrons repress resident R2 elements. As the uninserted:R2-inserted rDNA ratio drops, as it does in our deletion alleles, we observe a derepression of the R2 and a derepression of the X-linked arrays. The possibility that derepression of the X-linked rDNA array is controlled by both uninserted and R1/R2-inserted rDNA copy numbers can reconcile ours and Greil and Ahmad’s opposite observations. It is hard not to imagine that these events – uninserted rDNA expression and retrotransposon repression, both mediated by heterochromatin – are not related. The mechanism for dominance may be as simple as a translationally-limited production of some heterochromatin-critical protein. This simple model would also account for the known (Paredes and Maggert 2009b) and verified (Larson et al., 2012; Zhou et al., 2012) connection between rDNA copy number and suppression of heterochromatin-induced gene silencing. We failed to find evidence to support this hypothesis (Paredes, 2011), though the defect may be subtle enough, or broadly-distributed enough (Paredes et al. 2011), so as to avoid easy detection.

### Magnification of the *rDNA* – what is a “magnifying” condition?

rDNA magnification in Drosophila has a long and complex history. Three things stand out from this seminal and classic work. First, the rate of magnification is relatively low. Second, that not all chromosome combinations are subject to magnification. Third, that no mutant conditions were identified that increased the rate of magnification. What also stands out is that the conditions under which magnification occurred – bobbed alleles in combinations that heavily biased expression – match the conditions we find here to be those that disrupt a dominance relationship between X- and Y-linked rDNA arrays. Although the bulk of previous research was done on X-linked rDNA arrays in females, examples of Y-linked magnification were known. From our results, we predict that those magnifying Y chromosomes did not exhibit nucleolar dominance. In fact, we hypothesize that any condition that disrupts nucleolar dominance will induce magnification. In our experiments here, we showed that disruptions of chromosome pairing (e.g., SMC1 and CTCF) did not induce magnification appreciably, however defects in heterochromatin-induced gene silencing did. Our results are consistent with Greil and Ahmad, who showed that Su(var)3-9 mutants disrupted dominance. They also found that “tempering” a chromosome by longtime exposure to Su(var)3-9 mutation altered the chromosomes so that they persistently lacked dominance even in the absence of the Su(var)3-9 mutation. We do not yet know what change to the Y chromosome was induced by tempering, but we note that those tempered chromosomes have an altered uninserted:R2-inserted ratio, as our hypothesis predicts.

This hypothesis also predicts a coincidence of magnification and R2 expression. Simply stated, losing uninserted rDNA alters the uninserted:R2 ratio (derepressing the R2 elements) and derepresses the heterologous rDNA array (inducing magnification); these phenomena usually overlap, but do not necessarily. The coincidence of R2 expression and magnification has been noted before, though they had concluded that R2 expression was not alone sufficient for magnification. However, R2 expression has recently been proposed to be causal in magnification (Nelson et al. 2023). Is it, or is it not? Perhaps the explanation is not R2 expression per se, but rather the effect of R2 expression on derepression of the X-linked rDNA. In recent studies, altering R2 expression (by siRNA-dependent heterochromatin formation) may repress the rDNA arrays, and thus alter the dominance relationship. Much mechanistic work remains to be done, however our results suggest a unification of previously-disparate properties of the rDNA. Repression of the retrotransposons within the rDNA is desirable, and the mechanism by which it occurs causes a nucleolar dominance relationship. Disruption of this condition frees retrotransposon expression, induces retrotransposition, causes DNA damage, and results in rDNA magnification.

## Data Availability

Strains and plasmids are available upon request. The authors affirm that all data necessary for confirming the conclusions of the article are present within the article, figures, and tables.

## Acknowledgements

This work was completed in partial fulfillment of a Ph.D. in Genetics from the Genetics Graduate Interdisciplinary Program at the University of Arizona. S. Kindelay thanks Drs. Frans Tax, Nathan Ellis, and Bonnie Hurwitz for their advising, and Ergül Susamcı for advice, discussion, and support. Kami Ahmad shared strains used in his work, which were invaluable in ours.

## Funding

We gratefully acknowledge financial support from the National Institutes of Health to the laboratory (R01GM123640), the University of Arizona Cancer Center (P30CA023074), and the Bloomington Drosophila Stock Center (P40OD018537), as well as from the University of Indiana to the last. We also acknowledge generous resource support from the University of Arizona (for the University of Arizona Genetics Core). This work was further supported by a seed grant from the Department of Molecular & Cellular Biology. S. Kindelay is a recipient of the Initiative for Maximizing Student Diversity (IMSD) fellowship from the Graduate College at the University of Arizona, and of a Sloan Indigenous Foundation fellowship.

## Conflict of Interest

The authors declare they have no competing interests. Conflict of Interest certification is on file at the University of Arizona Office for Responsible Outside Interests.

---

Supplementary Figure 1. Further examples of variegation in the compound eyes and dorsal abdomens of y^1^ w^67c23^/Y, 590 males. See also Figure 1C.

Supplemental Figure 2. Schemes for generating rDNA deletions, varying total times of heat shock induction of I-CreI over 2-5 days. Schemes were generally assessed by monitoring male-to-female ratio (“M/F”).

Supplemental Figure 3. Lethal-phase analysis of all rDNA deletions generated in this study. See also Figure 2D.

Supplemental Figure 4. Analysis of rDNA copy number to (A) length of secondary constriction (including representative images of secondary constructions (B)), and to fluorescence in situ hybridization fluorescence intensity of R2 (C) and 18S (D). Statistical tests were Student’s t-test, and all comparisons failed to find significant differences in populations. Units in (A), (C), and (D) are arbitrary units, but are consistent between chromosome preparations. In (A), measurements are of the Y-linked rDNA, and in (C) and (D), measurements are of both X- and Y-linked rDNA (“(X)” and “(Y),” respectively). In (C) and (D), although there is difference between chromosomes, the wide variation makes predictive measurements (i.e., inferring rDNA copy number from fluorescence) impossible.

Supplemental Figure 5. qPCR analysis of rDNA copy number on the C(1)DX chromosome used in this study (genotype is C(1)DX, y^1^ f^1^ bb^0^/0).

Supplemental Figure 6. Quantification of bobbed and bobbed^+^ offspring from C(1)DX mothers and one of three fathers – Y, 10B (a wild-type Y chromosome) (Maggert and Golic, 2005; Paredes and Maggert 2009), Y, 10B^484-bb-L^ (a bobbed Y chromosome) (Paredes and Maggert, 2009), and Y, 590.

Supplemental Figure 7. Fluorescence in situ hybridization of R1 and R2 sequences to the C(1)DX chromosome in combination with different Y chromosomes. See also Figure 5.

Supplemental Figure 8. Examples of the categories of nucleolar dominance as exhibited in a wild-type (y w/Y, 590) genotype. See also Figure 6.

Supplemental Figure 9. Quantification and representative images of nucleolar dominance using a chromosome first characterized in (Greil and Ahmad, 2012).

Supplemental Figure 10. Quantification of uninserted, R1-inserted and R2 inserted rDNA copy numbers on wild-type Y chromosomes Y, 10B (A), Y, 590 (B) and bobbed rDNA deletion allele, Y, 10B^183^ (C). Statistical tests are Student’s t-test.

Supplemental Figure 11. Genotypes and sources of the strains used in this study.

